# AI-Guided Discovery of Small Molecule LILRB4 (ILT3) Inhibitors Reprograms Microglia and Reduces Amyloid Pathology

**DOI:** 10.64898/2026.06.12.731845

**Authors:** Somaya A. Abdel-Rahman, Moustafa T. Gabr

## Abstract

The inhibitory microglial receptor LILRB4 (ILT3) suppresses amyloid-beta clearance in Alzheimer’s disease (AD) through ApoE-dependent signaling but remains undrugged by small molecules. Here, we report the AI-guided discovery of small molecule inhibitors that directly disrupt the LILRB4-ApoE interaction. Ultralarge-scale screening of ∼500 million compounds identified small molecules that bind LILRB4 with nanomolar affinity and competitively block ApoE engagement, as validated across orthogonal biophysical assays. Structural and mutational analyses define a tractable interdomain pocket that mediates ligand recognition. In human iPSC-derived microglia, LILRB4 inhibition suppresses SHP1/2-dependent signaling, attenuates NF-κB activation, and restores Aβ uptake. The lead compound exhibits favorable pharmacokinetics with robust brain penetration and, upon oral administration, improves cognitive performance, reduces amyloid burden, and dampens neuroinflammation in the 5xFAD mouse model. These findings establish LILRB4 as a druggable neuroimmune checkpoint and demonstrate that small molecule disruption of inhibitory microglial signaling can restore disease-relevant function in vivo.

## INTRODUCTION

Alzheimer’s disease (AD) remains a major unmet clinical challenge, with disease-modifying therapies largely limited to amyloid-targeting antibodies that provide modest benefit and are constrained by limited brain penetration and safety concerns (1–5). Increasing evidence implicates microglia as central regulators of amyloid-β (Aβ) pathology, where disease progression reflects an imbalance between activating and inhibitory signaling pathways that control microglial function (2,6–10). Microglial responses are governed by a balance between activating receptors such as triggering receptor expressed on myeloid cells 2 (TREM2) and inhibitory receptors including CD33 and members of the leukocyte immunoglobulin-like receptor (LILR) family, which collectively shape phagocytic capacity and inflammatory signaling (4,5). Recent studies, including our own, have demonstrated that small molecule targeting of TREM2 can reprogram microglial function, supporting the feasibility of pharmacological modulation of neuroimmune checkpoints (11–14).

The inhibitory receptor, leukocyte immunoglobulin-like receptor B4 (LILRB4, ILT3), has recently emerged as a key microglial immune checkpoint that suppresses Aβ clearance through ApoE-dependent signaling (15–22). LILRB4 is highly upregulated in microglia from patients with AD, particularly in plaque-associated populations, where its expression correlates with ApoE levels and pathological markers including phosphorylated tau (23). Mechanistically, LILRB4 engagement activates immunoreceptor tyrosine-based inhibitory motifs (ITIMs), recruiting SHP1/2 phosphatases and suppressing cytoskeletal remodeling and phagocytic pathways required for effective plaque clearance. Antibody-mediated blockade of LILRB4 restores microglial activation, enhances Aβ engulfment, and reduces amyloid burden by approximately 50% in preclinical models, accompanied by suppression of interferon-response programs and reorganization of plaque-associated microglia (23). These findings establish LILRB4 as a compelling therapeutic target for AD and identify the LILRB4-ApoE interaction as a functionally defined interface amenable to pharmacological targeting.

Recent advances in artificial intelligence (AI) are transforming structure-based drug discovery, particularly for challenging targets lacking well-defined binding pockets (24–30). Among these, contrastive learning-based frameworks have introduced a shift from traditional affinity prediction to representation-based retrieval, enabling direct alignment of protein pockets and small molecules within a shared embedding space. DrugCLIP (28) exemplifies this approach by reframing virtual screening as a dense retrieval problem, where binding is inferred through embedding similarity rather than explicit scoring functions. This strategy mitigates the limitations of noisy affinity labels and enables ultralarge-scale screening across hundreds of millions of compounds with orders-of-magnitude improvements in computational efficiency relative to docking-based methods. Such approaches provide a powerful route to address protein–protein interaction interfaces that have historically been considered undruggable.

Here, we report a contrastive learning-based discovery strategy that establishes LILRB4 (ILT3) as a tractable small molecule target. Using DrugCLIP (28), which reframes virtual screening as a dense retrieval problem, we performed ultralarge-scale screening to identify candidate LILRB4 small molecule ligands. Biophysical validation and functional characterization in human microglia revealed inhibitors that restore phagocytic activity and modulate inhibitory signaling. Pharmacokinetic (PK) and in vivo studies further demonstrate central exposure and improved cognitive performance, along with reduced amyloid pathology and neuroinflammation in the 5xFAD mouse model. Together, this work provides a framework for small molecule modulation of microglial checkpoints and highlights retrieval-based AI screening as an effective strategy for targeting neuroimmune receptors.

## RESULTS

### Identification of a Ligandable Surface on LILRB4 for AI-guided Screening

To evaluate the potential of LILRB4 for small molecule modulation, we first sought to identify ligandable regions on its extracellular domain. Structural analysis of the human LILRB4 ectodomain (PDB ID: 3P2T) using PrankWeb (31) revealed a single, well-defined surface pocket located at the D1-D2 interdomain interface (Figure 1A). This region forms a contiguous, solvent-accessible cavity enriched in both hydrophobic and polar residues, including Leu132, Lys134, Leu144, Pro158, Met159, Val165, His166, and Tyr170. A closer structural inspection highlights the spatial arrangement of these residues along the interdomain groove, delineating a chemically heterogeneous binding environment consistent with protein-protein interaction (PPI) interfaces (Figure 1B).

**Figure 1.**
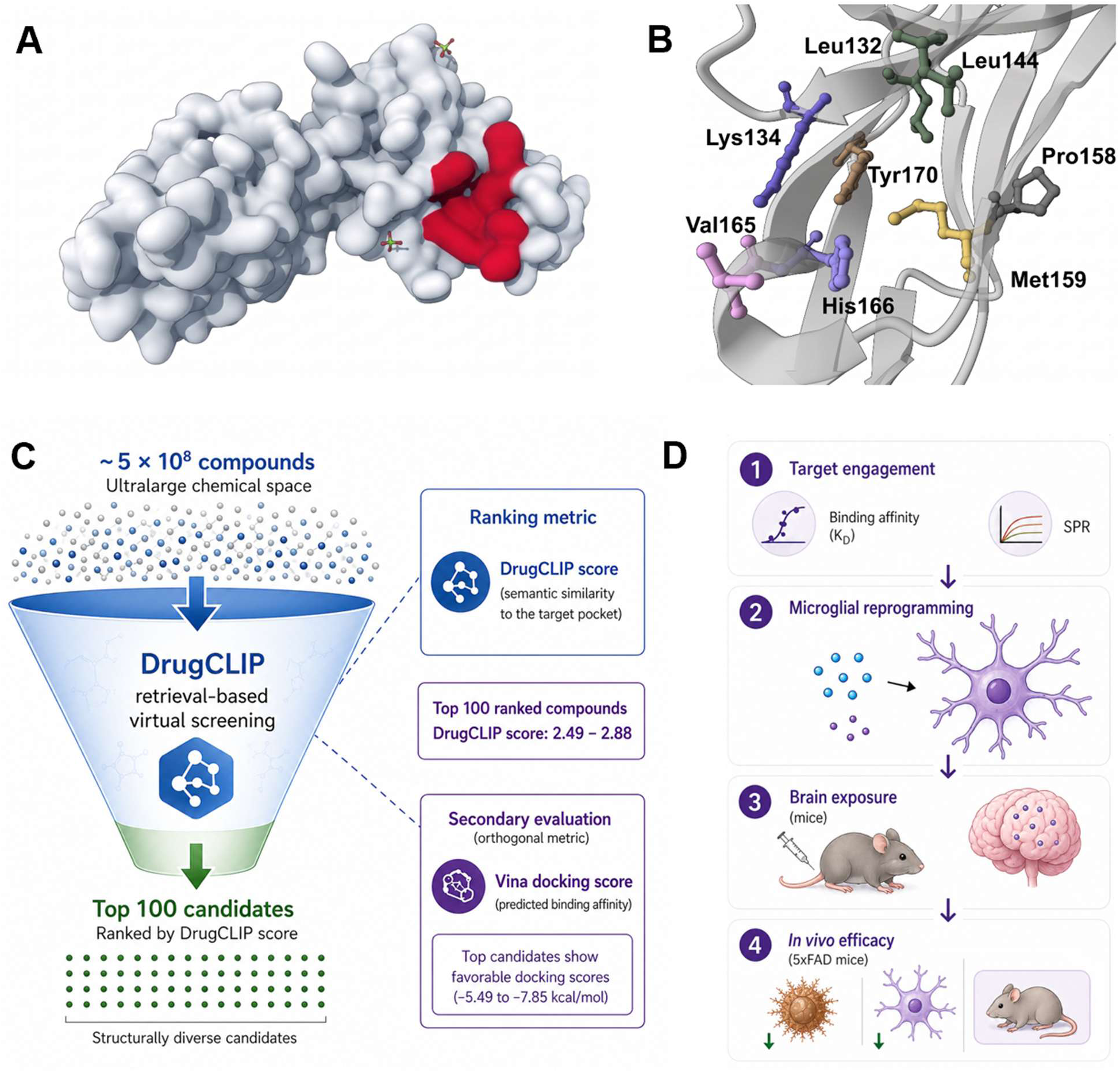
AI-guided discovery and validation of small molecule modulators targeting LILRB4. **(A)** Surface representation of the LILRB4 extracellular domain highlighting the predicted ligand-binding pocket (red) by PrankWeb. **(B)** Close-up view of the binding site with key residues lining the pocket, indicating potential interaction hotspots for small molecule engagement. **(C)** DrugCLIP-based virtual screening workflow. Approximately 5 × 10^8^ compounds were screened within an ultralarge chemical space using a retrieval-based approach. Compounds were ranked by DrugCLIP score (semantic similarity to the target pocket), and the top 100 candidates (score range: 2.49-2.88) were selected. Secondary evaluation using molecular docking (Vina) confirmed favorable predicted binding affinities (−5.49 to −7.85 kcal/mol), yielding structurally diverse candidates. **(D)** Experimental validation pipeline. Top candidates were evaluated for target engagement, functional modulation of microglial activity, brain exposure in mice, and in vivo efficacy in a 5xFAD mouse model.

Although the precise structural determinants of ApoE binding to LILRB4 remain incompletely defined, prior studies have established that LILRB4 functions as an inhibitory receptor that suppresses immune activation through ApoE-dependent signaling. Antibody-mediated disruption of this interaction restores immune function and attenuates disease-relevant signaling pathways, underscoring the therapeutic relevance of targeting this axis (32). Structural characterization of the blocking antibody h128-3/LILRB4 complex (PDB ID: 6K7O) further demonstrated that engagement of surface-exposed regions within the D1 domain through extensive hydrogen bonding interactions is sufficient to inhibit ApoE-dependent signaling. These findings suggest that modulation of LILRB4 does not require engagement of a deeply buried pocket but can instead be achieved through perturbation of accessible extracellular surfaces.

Guided by these insights, we selected the PrankWeb-identified interdomain pocket (Figure 1A) as the input site for large-scale virtual screening using DrugCLIP. As outlined in Figure 1C, approximately 5 × 10^8^ compounds spanning an ultralarge chemical space were evaluated using a retrieval-based framework that prioritizes candidates based on semantic similarity to the target binding environment. Compounds were ranked according to DrugCLIP score, and the top 100 candidates (score range: 2.49-2.88) were selected for further analysis. Detailed information for these candidates, including molecular identifiers, DrugCLIP scores, predicted docking energies (Vina), and SMILES representations, is provided in Table S1. Secondary evaluation using molecular docking (Autodock Vina) revealed consistently favorable predicted binding affinities (−5.49 to −7.85 kcal/mol), supporting the compatibility of these candidates with the identified pocket. Finally, the overall discovery and validation strategy is summarized in Figure 1D, highlighting the progression from in silico prioritization to experimental validation. Top-ranked compounds were subsequently advanced for biochemical target engagement, functional evaluation, and in vivo characterization, establishing a systematic pipeline for translating computational predictions into biologically relevant outcomes.

### Biophysical screening and validation of DrugCLIP-prioritized compounds

To experimentally validate the top-ranked candidates identified through DrugCLIP screening (Figure 1C; Table S1), we performed biophysical binding assays against the LILRB4 extracellular domain using microscale thermophoresis (MST), a solution-phase technique that detects ligand binding through changes in thermophoretic mobility of a fluorescently labeled protein.

All 100 compounds were first evaluated in a single-dose MST primary screen at a fixed concentration using the Dianthus platform. Binding responses were quantified using normalized fluorescence (F_norm_) relative to DMSO controls. Compounds exhibiting a ΔF_norm_ ≥ 5 units with consistent responses across replicates (n = 5) were classified as preliminary hits. Based on these criteria, 21 compounds were identified as initial hits (Figure 2A), corresponding to a hit rate of 21%. The distribution of F_norm_ values revealed a clear separation between binders and non-binders, with the majority of compounds clustering tightly around baseline and a subset displaying pronounced positive shifts, supporting the robustness and discriminatory power of the screening conditions. To benchmark assay performance and define the dynamic range, the anti-LILRB4 blocking antibody h128-3 was included as a positive control and exhibited high-affinity binding with a dissociation constant (Kd) of 2.51 nM (95%CI: 2.33-2.75 nM), confirming the sensitivity of the MST platform to detect strong interactions.

**Figure 2.**
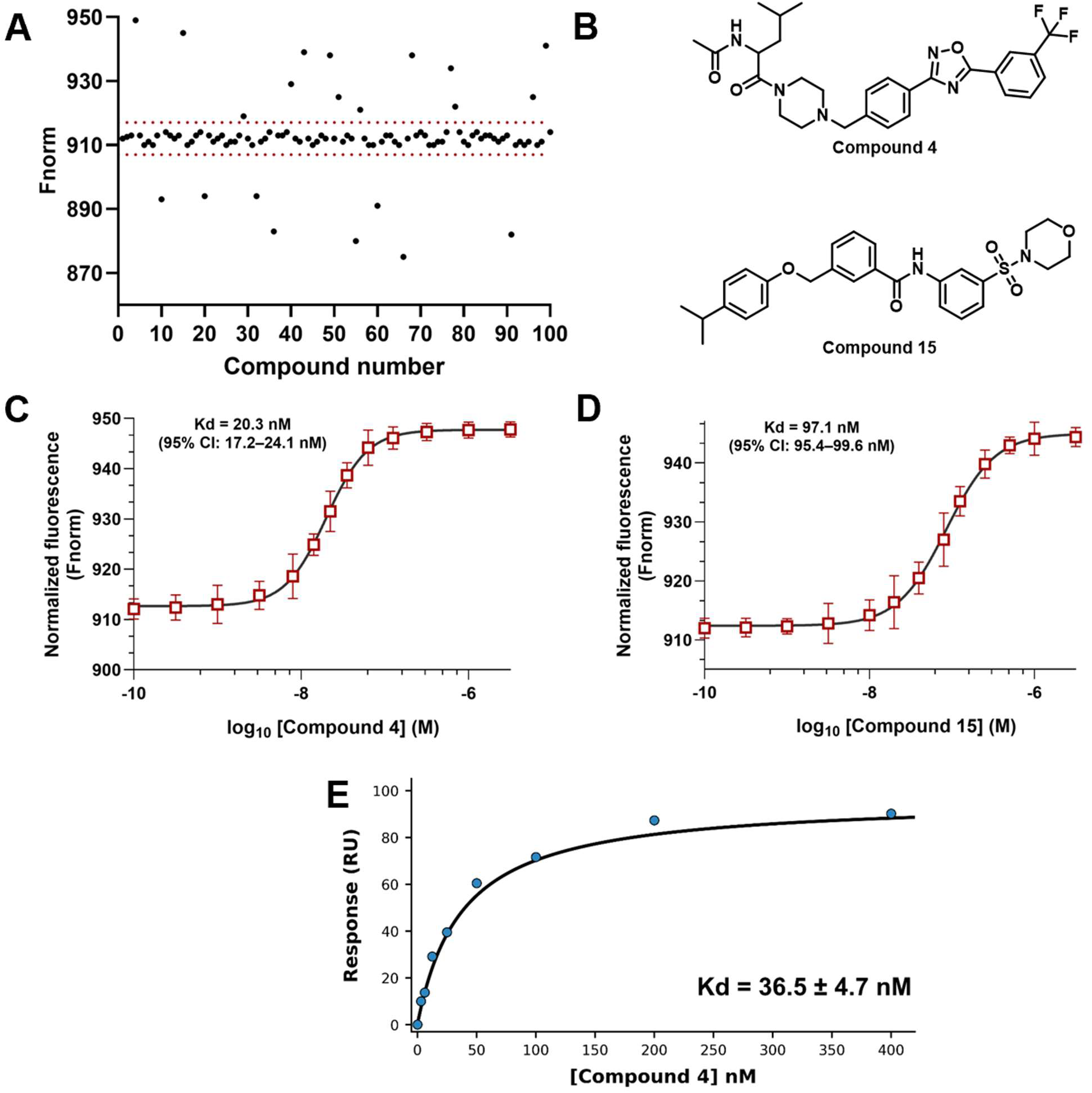
Experimental validation of DrugCLIP-prioritized compounds targeting LILRB4. **(A)** Single-dose MST screening of 100 DrugCLIP-selected compounds against the LILRB4 extracellular domain. Binding responses are shown as normalized fluorescence (F_norm_). Red dashed lines indicate the hit threshold (ΔF_norm_ ≥ 5). Compounds exceeding this threshold were classified as preliminary hits. **(B)** Chemical structures of the top two hits (compounds **4** and **15**) selected based on LILRB4 binding affinity. **(C)** Dose-dependent MST binding curve for compound **4**, showing a robust, saturable interaction with LILRB4 (Kd = 20.3 nM; 95% CI: 17.2-24.1 nM). **(D)** Dose-dependent MST binding curve for compound **15**, demonstrating a weaker but reproducible interaction (Kd = 97.1 nM; 95% CI: 95.4-99.6 nM). Data are shown as mean ± SD (n = 5). **(E)** Orthogonal validation of compound **4** binding by surface plasmon resonance (SPR), showing a concentration-dependent response with a hyperbolic binding profile (Kd = 36.5 ± 4.7 nM, n=3).

To further validate the primary hits, all 21 compounds were subjected to dose-dependent MST analysis. Among these, nine compounds (**4**, **15**, **20**, **32**, **49**, **55**, **56**, **66**, and **78**) exhibited concentration-dependent binding curves with well-defined saturation behavior, supporting their classification as validated LILRB4 binders. The remaining compounds showed weak, non-saturating, or inconsistent responses and were excluded from further analysis. The MST Kd values of the validated LILRB4 binders are provided in Table S2.

From this subset, two hits (compounds **4** and **15**, Figure 2B) emerged as the most potent small molecule LILRB4 binders from this study. MST binding curves for these compounds demonstrate robust and reproducible dose-dependent interactions with LILRB4, yielding Kd values of 20.3 nM (95% CI: 17.2-24.1 nM) for compound **4** and 97.1 nM (95% CI: 95.4-99.6) for compound **15** (Figures 2C and 2D). To confirm binding using an orthogonal technique, compound **4** was further evaluated by surface plasmon resonance (SPR). This analysis revealed a concentration-dependent increase in response units with a characteristic hyperbolic binding profile, yielding a Kd of 36.5 ± 4.7 nM (Figure 2E). The close agreement between MST and SPR measurements, despite differences in assay format and immobilization, reinforces the robustness of the observed compound **4**/LILRB4 interaction. Together, these results establish a robust and scalable workflow integrating single-dose MST screening (Figure 2A), quantitative dose-response validation (Figures 2C,D), and orthogonal SPR confirmation (Figure 2E), enabling efficient identification and prioritization of high-affinity small molecule binders targeting LILRB4.

### Structural and mutational validation of the LILRB4 binding pocket

To gain mechanistic insight into the binding mode of the top DrugCLIP-prioritized compounds, we performed structure-guided modeling followed by experimental validation using site-directed mutagenesis. Predicted 3D binding poses (from AutoDock Vina) revealed that both compounds **4** and **15** occupy a well-defined interdomain pocket within the LILRB4 extracellular domain (Figures 3A and 3D). In both cases, the ligands adopt conformations that maximize surface complementarity within a predominantly hydrophobic cavity, while forming discrete polar interactions with surrounding residues. Detailed 2D interaction mapping further highlighted the key residue-level contacts mediating ligand engagement (Figures 3B and 3E). For compound **4**, interactions were predicted with residues including Lys134, Leu144, Met159, Val165, and Thr163, involving a combination of hydrogen bonding and hydrophobic contacts. Compound **15** exhibited a partially overlapping interaction network within the same pocket, with prominent contributions from Lys134 and Leu144, alongside additional scaffold-specific contacts. These results suggest that while both compounds target a common binding site, they engage the pocket through distinct but related interaction patterns.

**Figure 3.**
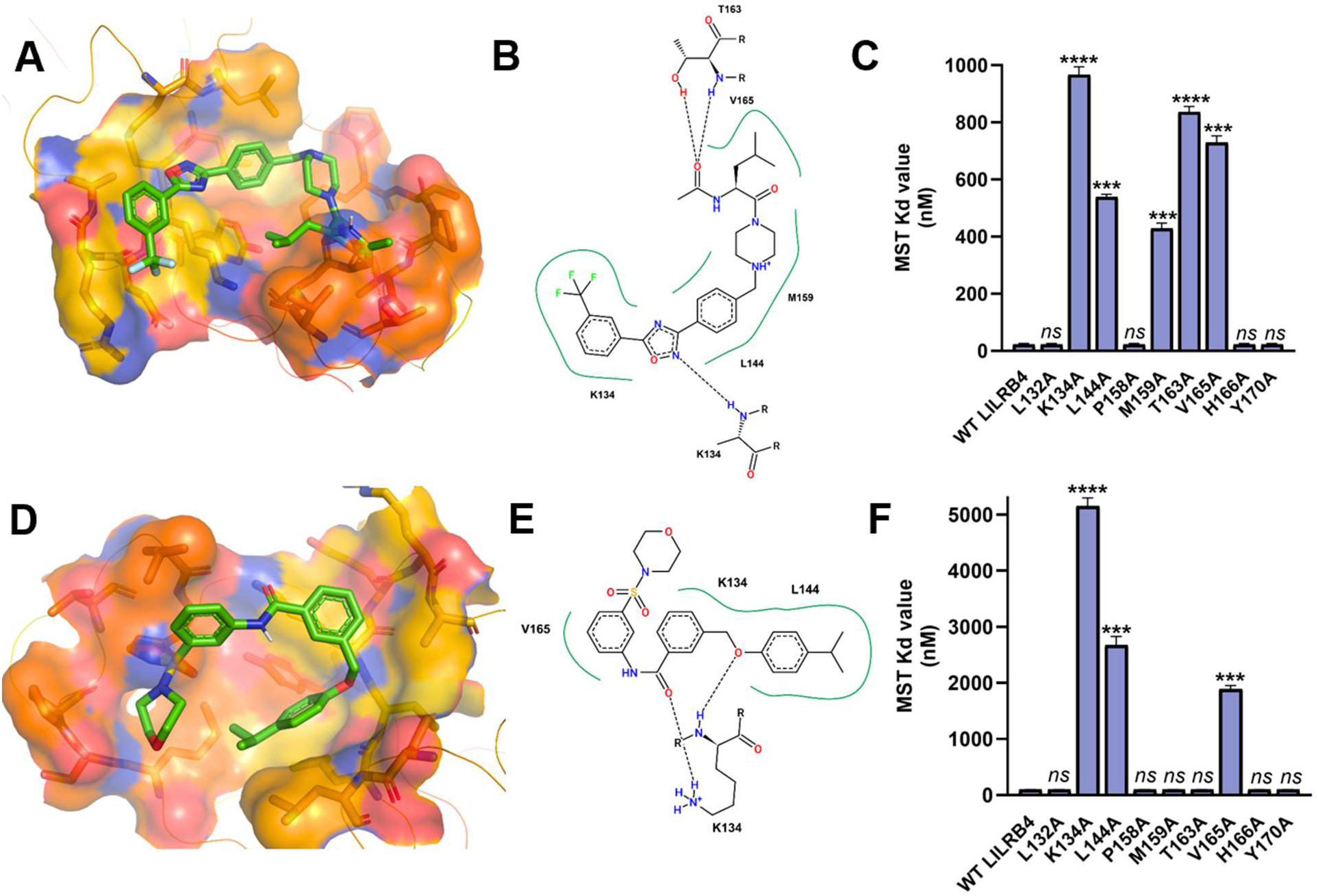
Structural and mutational validation of LILRB4 small molecule binding interactions. **(A)** Predicted 3D binding mode of compound **4** within the LILRB4 extracellular domain, highlighting the interdomain pocket and key interacting residues. The protein surface is shown with electrostatic coloring, and the ligand is depicted in stick representation. **(B)** 2D interaction map of compound **4** illustrating key contacts with residues including Lys134, Leu144, Met159, Val165, and Thr163. Hydrogen bonds and hydrophobic interactions are indicated by dashed lines. **(C)** Alanine scanning mutagenesis of predicted binding site residues and corresponding MST-derived binding affinities. Mutations at key positions (e.g., K134A, T163A, V165A) resulted in substantial loss of binding affinity, confirming their critical role in ligand engagement. Data are shown as mean ± SD (n = 5). Statistical significance is indicated relative to WT LILRB4. **(D)** Predicted 3D binding mode of compound **15** within the same LILRB4 pocket, showing a similar binding orientation with distinct interaction features. **(E)** 2D interaction map of compound **15** highlighting key residue contacts, including Lys134 and Leu144, supporting a shared binding region with compound-specific interactions. **(F)** MST-based mutational analysis for compound **15**, demonstrating residue-dependent effects on binding affinity. Key mutations (e.g., K134A and L144A) markedly disrupt binding, while others have minimal impact, indicating both shared and distinct interaction determinants compared to compound **4**. Data are shown as mean ± SD (n = 5).

To experimentally validate these predicted interactions, we performed alanine scanning mutagenesis of key residues (Leu132, Lys134, Leu144, Pro158, Met159, Thr163, Val165, His166, and Tyr170), followed by MST-based binding analysis. For compound **4**, mutation of several residues resulted in pronounced loss of binding affinity (Figure 3C). In particular, K134A, T163A, and V165A substitutions led to substantial increases in MST Kd values, indicating that these residues play critical roles in stabilizing ligand binding. Additional moderate effects were observed for L144A and M159A, consistent with their involvement in hydrophobic packing within the pocket. In contrast, mutations such as L132A, P158A, H166A, and Y170A had minimal impact, suggesting a more peripheral contribution to ligand recognition.

A similar but distinct pattern was observed for compound **15** (Figure 3F). The K134A mutation resulted in a dramatic loss of LILRB4 binding affinity, highlighting Lys134 as a central anchoring residue for both compounds. L144A also significantly disrupted binding, while V165A exhibited a compound-dependent effect, consistent with differences in ligand orientation and interaction geometry. Notably, several mutations that strongly affected compound **4** had reduced or negligible impact on compound **15**, reinforcing the notion of shared binding architecture with compound-specific interaction profiles.

Together, these data provide strong experimental support for the predicted binding pocket and establish a clear structure-function relationship governing ligand recognition. The convergence of 3D structural modeling (Figures 3A and 3D), interaction mapping (Figures 3B and 3E), and mutational validation (Figures 3C and 3F) demonstrates that DrugCLIP-derived compounds engage a defined and targetable site on LILRB4. Importantly, the identification of key residues such as Lys134, Leu144, Thr163, and Val165 as critical determinants of binding highlights potential hotspots for future structure-based optimization. These findings not only validate the accuracy of the DrugCLIP prioritization and docking framework but also establish a mechanistic foundation for rational ligand design targeting LILRB4. The ability of structurally distinct compounds to engage the same pocket through partially overlapping interaction networks further suggests that this site is both chemically tractable and adaptable, supporting its potential as a druggable interface for small molecule modulation.

### Inhibition of the LILRB4-ApoE interaction by DrugCLIP-derived compounds

Having established direct binding of DrugCLIP-prioritized compounds to a defined pocket on LILRB4 (Figure 3), we next asked whether these interactions translate into functional disruption of ligand binding. Given the reported interaction between LILRB4 and ApoE, we evaluated the ability of the top compounds to inhibit this PPI using complementary biochemical assays (Figure 4A). In a plate-based ELISA format, immobilized LILRB4 extracellular domain was incubated with ApoE in the presence of increasing concentrations of compounds **4** and **15**. Both compounds exhibited clear, dose-dependent inhibition of ApoE binding, yielding well-defined sigmoidal dose-response curves. Compound **4** showed potent activity with an IC_50_ of 40.6 nM (95%CI: 35.1-44.9 nM, Figure 4B), achieving near-complete inhibition at higher concentrations, while compound **15** displayed reduced potency with an IC_50_ value of 305.7 nM (95%CI: 241.2-374.5 nM, Figure S1).

**Figure 4.**
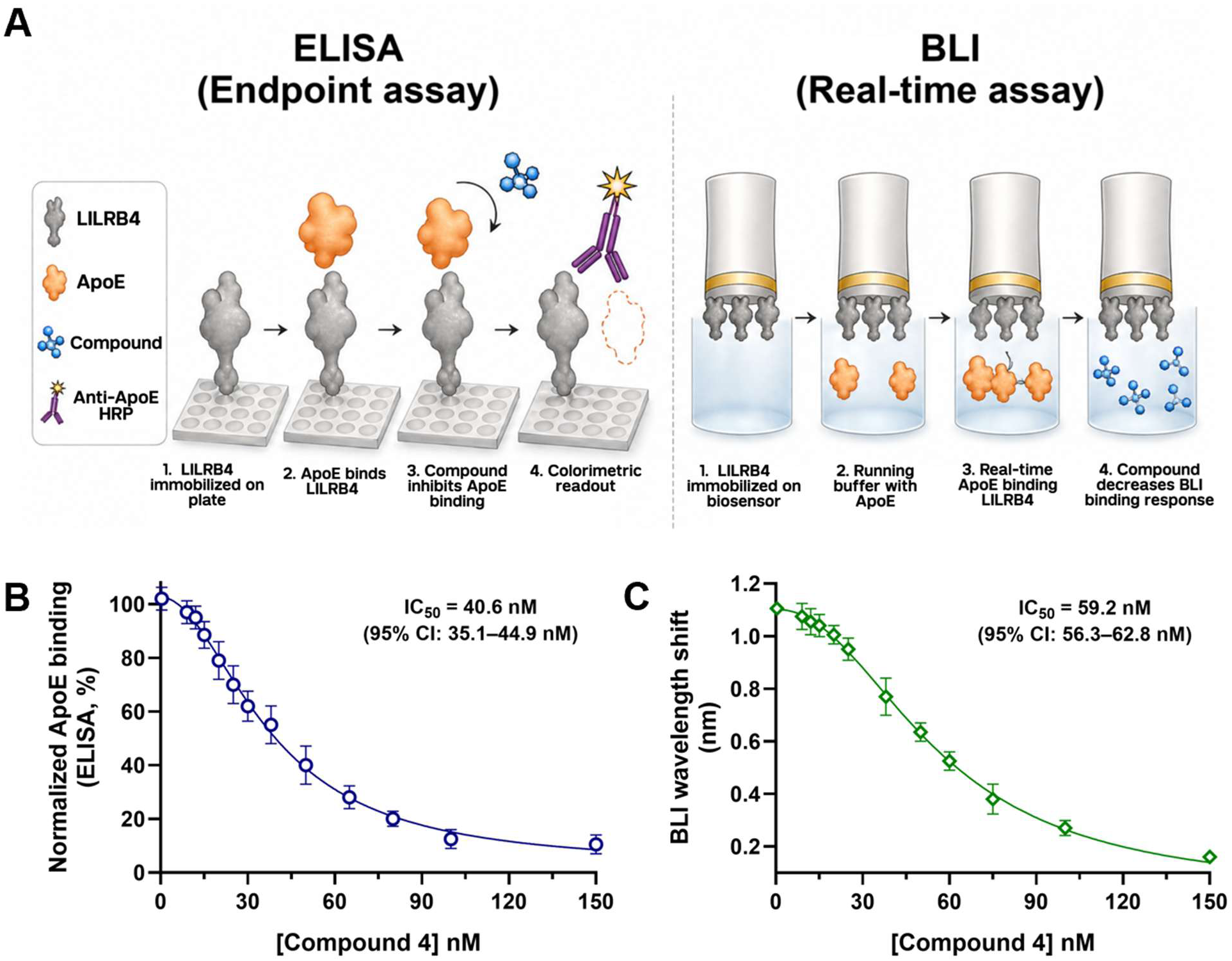
Orthogonal validation of compound 4 inhibition of the LILRB4-ApoE interaction. **(A)** Schematic of the ELISA (endpoint) and BLI (real-time) assays used to evaluate disruption of the LILRB4-ApoE interaction. In the ELISA format, LILRB4 is immobilized on a plate, ApoE binding is detected using an HRP-conjugated antibody, and compound-mediated inhibition is quantified by a decrease in colorimetric signal. In the BLI assay, LILRB4 is immobilized on a biosensor, ApoE association is monitored in real time, and compound binding reduces the wavelength shift corresponding to decreased complex formation. **(B)** Dose-dependent inhibition of ApoE binding to LILRB4 by compound **4** measured by ELISA. Data are presented as normalized ApoE binding (%). Data are shown as mean ± SD (n = 5). **(C)** BLI-based measurement of compound **4** activity for LILRB4-ApoE inhibition showing a concentration-dependent decrease in binding response. Data are shown as mean ± SD (n = 5).

To validate these findings using an orthogonal approach, we performed competitive binding experiments by biolayer interferometry (BLI), enabling real-time monitoring of ApoE association with immobilized LILRB4. Consistent with the ELISA results, compound **4** produced a concentration-dependent reduction in binding response with an IC_50_ value of 59.2 nM (95%CI: 56.3-62.8, Figure 4C), while compound **15** exhibited reduced potency with an IC_50_ value of 367.4 nM (95%CI: 334.1-395.6, Figure S2). The close agreement between ELISA and BLI across independent assay formats supports a direct and competitive disruption of the LILRB4-ApoE interaction.

Notably, these functional data align with the structural and mutational analyses (Figure 3), which place the compound binding site within or proximal to the predicted ApoE interaction interface. This spatial overlap supports a mechanism in which compound binding interferes with ligand engagement, either through direct competition or local perturbation of the binding surface. Together, these results demonstrate that DrugCLIP-identified binders not only engage LILRB4 with nanomolar affinity but also translate this interaction into functional disruption of a biologically relevant PPI. The strong concordance between ELISA and BLI, combined with structural validation, establishes a clear link between binding and biochemical function and provides a foundation for subsequent evaluation of downstream cellular effects.

### Pharmacological disruption of LILRB4-ApoE signaling restores microglial function in human iPSC-derived microglia

To determine whether disruption of the LILRB4-ApoE interaction translates into functional modulation in disease-relevant systems, we evaluated compound **4** in iPSC-derived human microglia under Aβ-driven conditions. Cells were exposed to Aβ42 oligomers in the presence or absence of ApoE to model ligand-engaged LILRB4 signaling, followed by treatment with compound **4** (50, 250, and 500 nM).

We first assessed proximal signaling by quantifying phosphorylation of SHP1 and SHP2, key phosphatases recruited to LILRB4 ITIM motifs. Co-stimulation with Aβ42 and ApoE induced a marked increase in phospho-SHP1 and phospho-SHP2 relative to both vehicle and Aβ alone (Figure 5A,B), consistent with activation of LILRB4-dependent inhibitory signaling. Treatment with compound **4** resulted in a clear, dose-dependent reduction in SHP1 and SHP2 phosphorylation, with partial suppression at 50 nM and substantial inhibition at 250 and 500 nM (Figure 5A,B), indicating effective blockade of ligand-induced receptor signaling in a cellular context.

**Figure 5.**
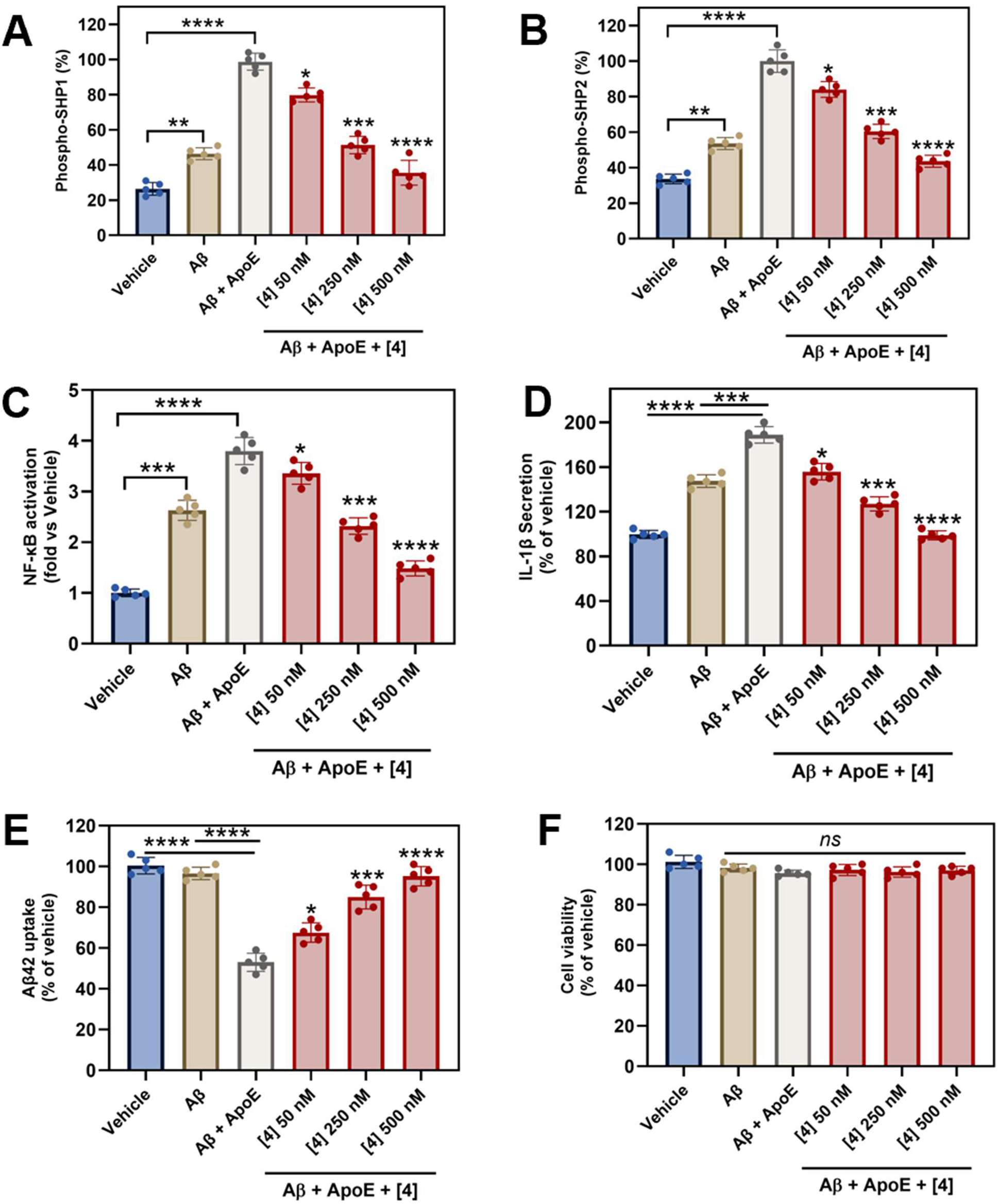
Compound 4 suppresses LILRB4-dependent signaling, attenuates neuroinflammation, restores amyloid-β uptake, and shows no cytotoxicity in iPSC-derived human microglia. **(A,B)** Phosphorylation of SHP1 **(A)** and SHP2 **(B)** following stimulation with Aβ and ApoE. Co-treatment with ApoE markedly enhanced SHP1/2 phosphorylation relative to Aβ alone, consistent with activation of LILRB4 signaling. Compound **4** reduced phospho-SHP1 and phospho-SHP2 levels in a dose-dependent manner. **(C)** NF-κB activation (fold vs vehicle). Aβ induced NF-κB activation, which was further increased by ApoE. Compound 4 suppressed NF-κB activation in a dose-dependent manner. **(D)** IL-1β secretion (% of vehicle). Aβ increased IL-1β release, enhanced by ApoE. Compound **4** reduced IL-1β in a dose-dependent manner. **(E)** Aβ42 uptake (% of vehicle). Aβ alone did not significantly alter uptake, whereas co-treatment with ApoE markedly impaired microglial Aβ uptake. Compound **4** restored Aβ uptake in a dose-dependent manner. **(F)** Cell viability (% of vehicle). No significant changes in viability were observed across all conditions, indicating that the effects of Compound 4 are not due to cytotoxicity. Data are presented as mean ± SD (n = 5). Statistical significance was determined by one-way ANOVA with appropriate multiple-comparisons tests. Comparisons between Aβ and vehicle, and between Aβ + ApoE and control conditions, are indicated where relevant. For compound-treated groups, statistical comparisons were performed relative to the Aβ + ApoE condition unless otherwise indicated. \*\**p* < 0.01, \*\*\**p* < 0.001, \*\*\*\**p* < 0.0001, and *ns* denotes non-significant.

We next examined downstream inflammatory signaling. NF-κB activation, quantified as fold-change relative to vehicle, was significantly elevated upon Aβ42 stimulation and further enhanced by ApoE co-treatment (Figure 5C). Compound **4** attenuated NF-κB activation in a concentration-dependent manner, reducing activity toward baseline levels at higher concentrations (Figure 5C). The concordance between reduced SHP1/2 phosphorylation (Figure 5A,B) and suppression of NF-κB signaling (Figure 5C) supports a mechanistic link between LILRB4 engagement and downstream transcriptional regulation of inflammatory pathways in microglia.

To assess functional consequences, we measured secretion of the pro-inflammatory cytokine IL-1β. Aβ42 exposure increased IL-1β release, which was further potentiated by ApoE (Figure 5D), consistent with an amplified inflammatory state. Compound **4** significantly reduced IL-1β secretion across all tested concentrations, with the strongest suppression observed at 250 and 500 nM, mirroring its effects on upstream signaling pathways. Given the central role of microglia in amyloid clearance, we evaluated Aβ42 uptake as a disease-relevant functional readout. Aβ42 alone did not significantly alter uptake relative to vehicle, whereas co-treatment with ApoE markedly impaired Aβ internalization (Figure 5E), consistent with a dysfunctional microglial phenotype. Treatment with compound **4** restored Aβ uptake in a dose-dependent manner, with significant recovery observed at 250 and 500 nM (Figure 5E).

Importantly, no significant changes in cell viability were observed across all treatment conditions (Figure 5F), indicating that the observed modulation of signaling, inflammation, and uptake is not attributed to cytotoxic effects. Collectively, these results establish a direct functional link between ApoE-mediated LILRB4 activation and suppression of microglial signaling and effector functions. When considered alongside the biochemical and biophysical validation, these findings demonstrate that DrugCLIP-identified binders translate target engagement into functional rescue in human microglial models, supporting LILRB4 as a tractable target for modulating neuroimmune dysfunction in AD.

### Cross-species target engagement and PK profiling support in vivo evaluation in 5xFAD mice

Prior to in vivo efficacy studies, we sought to establish cross-species target engagement and characterize the PK properties of compound **4** to ensure adequate systemic and central nervous system (CNS) exposure. To assess cross-reactivity with murine LILRB4, we performed MST using the recombinant extracellular domain of mouse LILRB4. Compound **4** exhibited high-affinity binding with a Kd of 17.5 nM (95%CI 13.1-21.4 nM, Figure S3), comparable to that observed for the human protein, confirming conserved target engagement across species and supporting the use of murine models for in vivo evaluation.

To contextualize in vivo PK, key in vitro ADME properties of compound **4** were first assessed. In human and mouse liver microsomes, compound **4** exhibited half-lives (t_1/2_) of 58 min and 42 min, corresponding to intrinsic clearance values of 18.6 mL/min/kg and 31.4 mL/min/kg, respectively, consistent with the moderate systemic clearance observed in vivo. Human and mouse plasma stability studies showed 92.4% and 95.2% parent compound remaining after 2 h, indicating minimal degradation under assay conditions. Plasma protein binding was 87.5% (fᵤ,p = 0.125), while brain tissue binding yielded an unbound fraction (fᵤ,brain = 0.10), enabling estimation of free CNS exposure.

We next performed PK studies in C57BL/6 mice following single-dose administration. After intravenous (IV) dosing at 2 mg/kg, compound **4** displayed a plasma half-life (t_1/2_) of 2.8 h, systemic clearance (CL = 12.6 mL/min/kg), and a volume of distribution (V_d_ = 2.9 L/kg), corresponding to a plasma AUC_0-∞_ of 3.8 µM·h. Following oral (PO) administration at 10 mg/kg, compound 4 achieved a plasma C_max_ of 1.9 µM at T_max = 0.75 h, with an AUC_0-∞_ of 7.81 µM·h, yielding an oral bioavailability of ∼42%, consistent with the IV-PO exposure relationship.

Given the therapeutic objective of modulating neuroimmune signaling in the CNS, brain exposure was evaluated following oral dosing. Compound **4** reached a brain C_max_ of 1.1 µM at 1 h post-dose, with a brain AUC_0-∞_ of 5.76 µM·h and a brain-to-plasma exposure ratio (K_p_) of ∼0.7. Correction for unbound fractions yielded a K_p,uu_ of ∼0.55, indicating efficient equilibration of free drug across the blood-brain barrier. The quantitative agreement between systemic exposure and CNS penetration establishes a coherent pharmacokinetic framework consistent with the observed in vivo efficacy in the 5xFAD model.

### In vivo efficacy of compound 4 in the 5xFAD model of AD

Having established cross-species target engagement and favorable PK properties with robust brain penetration, we next evaluated the in vivo efficacy of compound **4** in the 5xFAD transgenic mouse model of AD, which exhibits early amyloid deposition, neuroinflammation, and cognitive impairment. The study design is summarized in Figure 6A, and PK profiling confirmed sustained plasma and brain exposure following oral administration of 10 mg/kg of compound **4** (Figure 6B).

**Figure 6.**
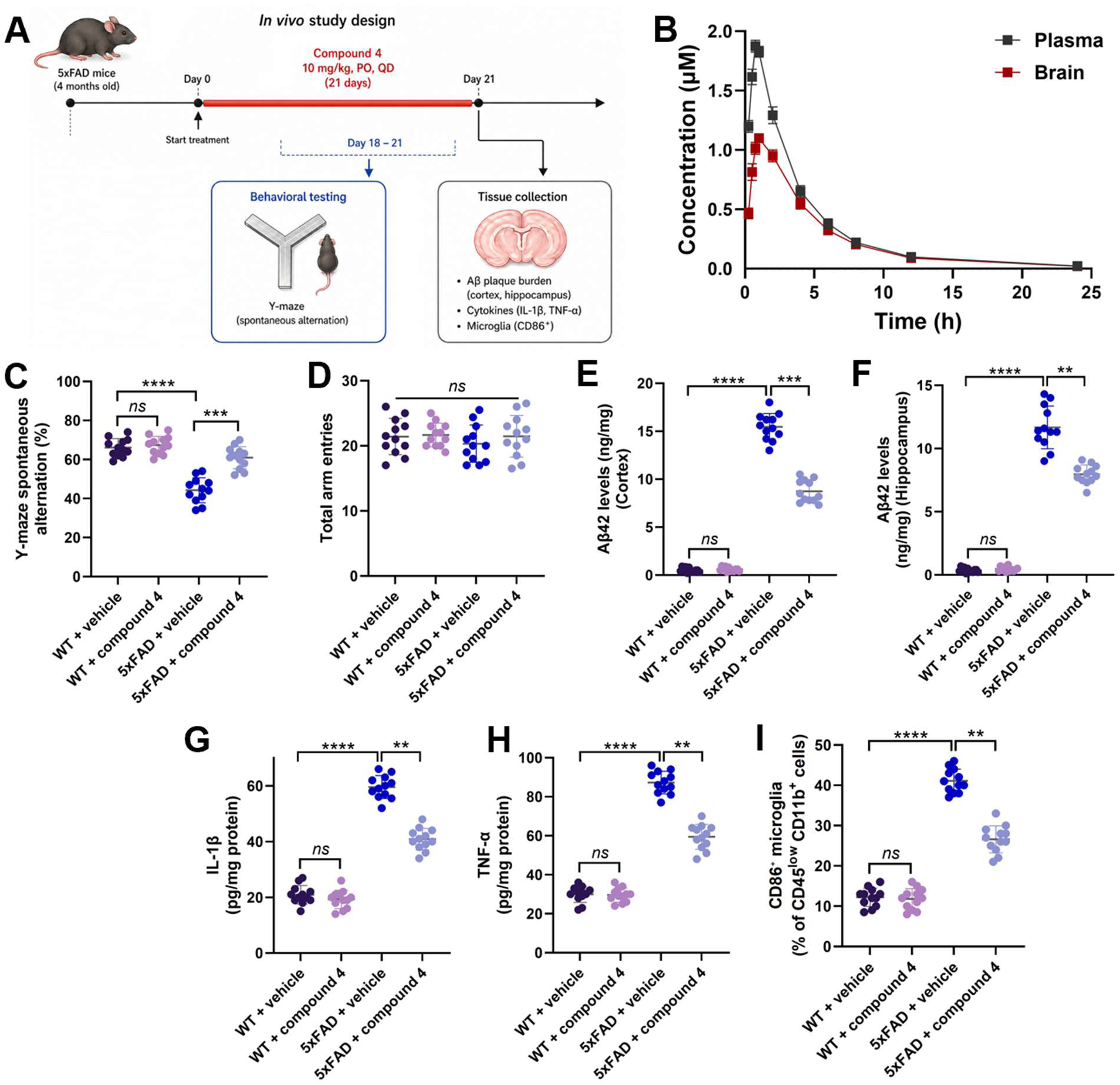
In vivo efficacy of compound 4 in 5xFAD mice. **(A)** Study design. Four-month-old 5xFAD mice were treated with compound **4** (10 mg/kg, PO, QD) for 21 days. Behavioral testing was performed during the final week, followed by tissue collection on Day 21. **(B)** Plasma and brain concentration-time profiles of compound **4** following oral administration, demonstrating sustained brain exposure. **(C)** Y-maze spontaneous alternation performance. 5xFAD mice exhibited impaired alternation compared to WT controls, which was significantly improved upon treatment with compound **4**. **(D)** Total arm entries in the Y-maze, showing no significant differences between groups, indicating no effect on locomotor activity. **(E, F)** Aβ42 levels in cortex **(E)** and hippocampus **(F)**. 5xFAD mice displayed elevated Aβ42 levels relative to WT mice, which were significantly reduced following compound **4** treatment. **(G, H)** Pro-inflammatory cytokine levels in brain tissue, including IL-1β **(G)** and TNF-α **(H)**, were elevated in 5xFAD mice and attenuated upon treatment. **(I)** CD86⁺ activated microglia (% of CD45^low^ CD11b^+^ microglia). 5xFAD mice showed increased microglial activation, which was reduced following treatment with compound **4**. Data are presented as mean ± SD (n = 12 per group). Statistical significance was determined by one-way ANOVA with appropriate post hoc multiple comparison tests. \*\**p* < 0.01, \*\*\**p* < 0.001, \*\*\*\**p* < 0.0001; *ns*, denotes non-significant.

Male and female 5xFAD mice and wild-type (WT) littermates (4 months of age) were randomized into treatment groups (n =12 per group; 6 males and 6 females) and administered compound **4** (10 mg/kg, PO, QD) or vehicle for 21 days (Figure 6A). Four-month-old 5xFAD mice were selected as this stage exhibits established amyloid pathology and neuroinflammation while retaining responsiveness to therapeutic intervention.^33^ Behavioral testing was conducted during the final week, followed by tissue collection for biochemical analyses. To assess functional outcomes, hippocampal-dependent working memory was evaluated using the Y-maze spontaneous alternation assay (Figure 6C). Vehicle-treated 5xFAD mice displayed a marked impairment relative to WT controls, consistent with cognitive deficits in this model. Treatment with compound **4** significantly improved alternation behavior in 5xFAD mice, restoring performance toward WT levels, while having no significant effect in WT animals (Figure 6C). Importantly, total arm entries were comparable across all groups (Figure 6D), indicating that the observed effects were not due to alterations in locomotor activity.

We next examined whether behavioral improvements were associated with modulation of amyloid pathology. Quantification of Aβ42 levels revealed a substantial increase in both cortex (Figure 6E) and hippocampus (Figure 6F) in 5xFAD mice relative to WT controls. Treatment with compound **4** significantly reduced Aβ42 levels in both regions, with ∼40% reduction in cortex and ∼35% reduction in hippocampus. Given the established link between amyloid pathology and neuroinflammation, we next assessed inflammatory signaling. Vehicle-treated 5xFAD mice exhibited elevated levels of pro-inflammatory cytokines, including IL-1β and TNF-α. Treatment with compound **4** significantly attenuated both cytokines (Figure 6G, H), indicating suppression of the inflammatory response in vivo.

To further evaluate microglial activation, we quantified CD86⁺ microglia as a marker of a pro-inflammatory phenotype by flow cytometry. 5xFAD mice showed a marked increase in CD86⁺ microglia compared to WT controls, whereas treatment with compound **4** significantly reduced this population (Figure 6I), consistent with a shift toward a less activated microglial state. Collectively, these data demonstrate that Compound **4** improves cognitive performance (Figure 6C), reduces amyloid burden (Figure 6E, F), and attenuates neuroinflammatory signaling (Figure 6G-I) in the 5xFAD model, without affecting general locomotor activity (Figure 6D), supporting its potential as a disease-modifying therapeutic targeting neuroimmune dysfunction in AD.

In summary, we demonstrate that LILRB4 can be directly targeted by small molecules to reverse inhibitory microglial signaling and restore disease-relevant function in AD models. These findings establish a new paradigm for pharmacological modulation of neuroimmune checkpoints and provide a foundation for the development of LILRB4-directed therapeutics. More broadly, this work illustrates how AI-guided discovery can unlock previously inaccessible targets, enabling new therapeutic strategies for complex neurodegenerative diseases.

## MATERIALS AND METHODS

### AI-guided screening for LILRB4-targeted small molecules

The crystal structure of the human LILRB4 extracellular domain (PDB ID: 3P2T) was retrieved from the Protein Data Bank. The structure was energy-minimized using the AMBER ff14SB force field to remove steric clashes while preserving backbone geometry. Potential ligandable pockets were identified using PrankWeb 4 (31), which employs a machine learning-based approach for binding site prediction. The highest-ranked pocket, located at the D1-D2 interdomain interface, was selected for downstream screening based on pocket score, solvent accessibility, and proximity to previously implicated functional regions.

An ultralarge virtual library (∼5 × 10⁸ compounds) was assembled from commercially accessible and enumerated chemical spaces (Enamine, ChemDiv, Vitas-M, and Princeton BioMolecular Research). All compounds were standardized using RDKit, including removal of salts, normalization of protonation states at pH 7.4, and generation of canonical SMILES. Virtual screening was performed using the DrugCLIP framework (28), a contrastive learning-based model that embeds protein pockets and small molecules into a shared latent space. The LILRB4 pocket was encoded and queried against the compound library using cosine similarity. Compounds were ranked based on DrugCLIP score (semantic similarity), and the top 100 candidates (score range: 2.49-2.88) were selected. Top-ranked compounds were subjected to molecular docking using AutoDock Vina. The docking grid was centered on the PrankWeb-identified pocket.

### Microscale thermophoresis (MST)

MST experiments were performed using the Dianthus NT.23 (NanoTemper Technologies). Purified His-tagged human LILRB4 protein was fluorescently labeled with RED-tris-NTA 2nd Generation dye (NanoTemper Technologies) according to the manufacturer’s instructions. Labeled protein (final concentration: 30 nM) was incubated with test compounds at a fixed concentration (10 µM) in MST buffer (PBS supplemented with 0.05% Tween-20 and 0.5% DMSO).

Fluorescence was measured in Dianthus 384-well plates (NanoTemper Technologies, Catalog# DI-P001A) with LED power set to 40%. Fluorescence was recorded at 670 nm and 650 nm, and normalized fluorescence (F_norm_) was calculated as the ratio of F670/F650. Binding responses were quantified as F_norm_. Compounds with ΔF_norm_ ≥ 5 units and reproducible responses across replicates (n = 5) were classified as hits. Dissociation constants (Kd) were obtained by fitting concentration-response curves using a four-parameter nonlinear regression model in GraphPad Prism 10. Reported values represent mean ± SD from at least three independent experiments.

### Surface plasmon resonance (SPR)

SPR validation of LILRB4 binding was performed using Biacore™ 8K SPR system (Cytiva, Marlborough, MA, USA). LILRB4-His Protein (10 μg/mL in PBS, pH 7.4) was immobilized on a Series S Sensor Chip CM5 (29104988, Cytiva, Marlborough, MA, USA) using a commercial amine coupling kit (BR100050, Cytiva, Marlborough, MA, USA). Immobilization was performed at a flow rate of 10 μL/min for 420 s, followed by blocking with ethanolamine. The flow cell with solely ethanolamine-block on the same channel served as the reference. Immobilization buffer: PBS-P+ (28995084, Cytiva, Marlborough, MA, USA). Briefly, gradient concentrations of the compound were prepared in the assay buffer, and injected over the sensor chip in a single-cycle kinetics model with a flow rate at 30 μL/min for 120 s per injection. After each injection, a 30 s-regeneration was performed at a flow rate of 30 μL/min using the regeneration buffer. Assays were performed in triplicate, and Kd values are reported as mean ± SD.

### ELISA-based competition assay

LILRB4 extracellular domain (2 µg/mL) was immobilized on 96-well high-binding plates (Corning) overnight at 4°C. Plates were blocked with 5% BSA in PBS for 1 h at room temperature. Recombinant human ApoE (R&D Systems) was added at a fixed concentration (100 nM) in the presence of serial dilutions of compounds. Bound ApoE was detected using an HRP-conjugated anti-ApoE antibody, followed by TMB substrate development. Absorbance was measured at 450 nm using a Tecan plate reader. Data were normalized to DMSO controls and fitted to determine IC_50_ values using nonlinear regression (GraphPad Prism v10). Reported values represent mean ± SD (n=5).

### Biolayer interferometry (BLI)

BLI experiments were performed using an Octet RED96 system (Sartorius). LILRB4 protein was immobilized onto Ni-NTA biosensors, followed by equilibration in assay buffer (PBS + 0.05% Tween-20). ApoE (100 nM) was associated in the presence of increasing concentrations of compounds. Binding responses were monitored as wavelength shifts, and inhibition curves were generated to determine IC_50_ values. Data were analyzed using Octet Data Analysis software. Reported values represent mean ± SD (n=5).

### Site-directed mutagenesis

Alanine substitutions (L132A, K134A, L144A, P158A, M159A, T163A, V165A, H166A, Y170A) were introduced into the human LILRB4 extracellular domain using the QuikChange Lightning Site-Directed Mutagenesis Kit (Agilent Technologies) according to the manufacturer’s protocol. Briefly, mutagenic primers were designed to incorporate the desired codon substitutions and PCR amplification was performed using a high-fidelity polymerase. Parental methylated DNA was digested with DpnI, and the resulting plasmids were transformed into *E. coli* DH5α cells. All mutations were confirmed by Sanger sequencing.

Recombinant wild-type and mutant LILRB4 proteins were expressed and purified as described above. Protein integrity and purity (>90%) were verified by SDS-PAGE and analytical size-exclusion chromatography. Binding affinities of mutant proteins were evaluated by MST under identical conditions to wild-type controls, and Kd values were obtained from nonlinear regression fits as described above.

### Human iPSC-derived microglia assays

Human iPSC-derived microglia (FujiFilm Cellular Dynamics) were cultured in microglia maintenance medium (Axol Bioscience Ltd, Catalog #ax0660) supplemented according to the supplier’s recommendations at 37 °C in a humidified 5% CO_2_ incubator. Cells were plated at 50,000 cells per well in poly-D-lysine-coated 96-well plates and allowed to recover for 24 h prior to treatment.

Aβ_42_ oligomers were prepared by dissolving lyophilized Aβ_42_ peptide (≥95% purity; AnaSpec, Catalog# AS-20276) in HFIP, followed by evaporation, resuspension in DMSO, and oligomerization in PBS at 4 °C for 24 h. Cells were treated with Aβ_42_ (1 μM) in the presence or absence of recombinant human ApoE (100 nM), followed by addition of compound **4** at indicated concentrations (50, 250, and 500 nM). Treatments were performed for 24 h. Vehicle controls contained equivalent DMSO concentrations (<0.5%).

### Signaling assays

Phosphorylation of SHP1 and SHP2 was quantified using phospho-specific ELISA kits (abcam, catalog# ab279924 and ab314344, respectively) following the manufacturer’s protocol. Briefly, cells were lysed in ice-cold lysis buffer supplemented with protease and phosphatase inhibitors. Lysates were normalized for total protein content (BCA assay), and equal amounts were loaded per well. Reported values represent mean ± SD (n=5).

NF-κB activation was measured using a luciferase reporter assay in human iPSC-derived microglia. Cells were plated at 50,000 cells per well in white 96-well plates and transfected with an NF-κB firefly luciferase reporter (Promega) and a Renilla luciferase control plasmid (Promega) using Lipofectamine 3000 (Thermo Fisher Scientific, Cat. #L3000008). Twenty-four hours post-transfection, cells were treated with Aβ42 oligomers (1 μM) ± ApoE (100 nM) and compound **4** (50-500 nM) for 24 h. Luciferase activity was measured using the Dual-Glo Luciferase Assay System (Promega, Cat. #E2920). NF-κB activity was calculated as the ratio of firefly to Renilla signal and expressed as fold change relative to vehicle controls. Reported values represent mean ± SD (n=5).

### Cytokine measurements

IL-1β levels in culture supernatants were quantified using a human IL-1β ELISA kit (R&D Systems, catalog# DLB50). Supernatants were collected, clarified by centrifugation (1,000 × g, 5 min), and analyzed according to the manufacturer’s instructions. Cytokine concentrations were interpolated from standard curves and normalized where appropriate. Reported values represent mean ± SD (n=5).

### Aβ uptake assay

Fluorescently labeled Aβ_42_ (HiLyte Fluor 555-conjugated from AnaSpec, catalog# AS-60480-01) was added to cells at a final concentration of 500 nM following treatment conditions described above. After incubation (4 h), cells were washed extensively with PBS and treated with trypan blue to quench extracellular fluorescence. Intracellular fluorescence was quantified using a Tecan Spark plate reader and normalized to cell number (total protein).

### Cell viability

Cell viability was assessed using the CellTiter-Glo luminescent assay (Promega, catalog#G7570) according to the manufacturer’s protocol. Following treatment, reagent was added directly to wells, incubated for 10 min at room temperature, and luminescence was measured. Viability was normalized to vehicle-treated controls. Reported values represent mean ± SD (n=5).

### PK studies

PK studies were conducted in male C57BL/6 mice (8-10 weeks old; n = 5 per time point). Compound **4** was administered by intravenous injection (2 mg/kg) or oral gavage (10 mg/kg) in a formulation consisting of 10% DMSO, 40% PEG400, and 50% sterile saline (v/v/v), prepared fresh prior to dosing. Blood samples were collected at 0.25, 0.5, 1, 2, 4, 8, and 24 h post-dose via tail vein sampling into EDTA-coated tubes. Plasma was isolated by centrifugation at 3,000 × g for 10 min at 4 °C.

For CNS exposure analysis, mice were perfused with ice-cold PBS before brain collection to minimize residual blood contamination. Brain tissues were harvested, weighed, and homogenized in ice-cold PBS. Compound concentrations in plasma and brain homogenates were quantified by LC-MS/MS following protein precipitation with acetonitrile containing an internal standard. Chromatographic separation was performed on a C18 column, and analytes were detected in positive electrospray ionization mode using multiple reaction monitoring. Calibration curves were prepared in blank mouse plasma or brain homogenate matrix and used to calculate compound concentrations. PK parameters were calculated by noncompartmental analysis using Phoenix WinNonlin. Brain-to-plasma exposure ratios were calculated from AUC values. Unbound brain-to-plasma ratios (K_p,uu_) were calculated using measured plasma and brain unbound fractions.

### In vivo efficacy in 5xFAD mice

All procedures were approved by the Institutional Animal Care and Use Committee (IACUC, Protocol #2024-0006). Male and female 5xFAD mice and wild-type littermates (4 months old) were randomized into treatment groups (n = 12 per group; 6 males and 6 females per group).

Compound **4** was administered by oral gavage at 10 mg/kg once daily (QD) for 21 days. Vehicle-treated animals received formulation only. Treatment allocation, behavioral testing, biochemical assays, flow cytometry analysis, and data quantification were performed by investigators blinded to genotype/treatment.

### Behavioral testing

Cognitive function was assessed using the Y-maze spontaneous alternation assay. Mice were placed in the maze and allowed to explore freely for 8 min. Alternation percentage was calculated as the ratio of sequential entries into all three arms over total possible alternations. Total arm entries were recorded to control for locomotor activity.

### Amyloid quantification

Cortex and hippocampus were microdissected separately, snap-frozen, and homogenized in ice-cold extraction buffer containing protease inhibitors. Tissue homogenates were centrifuged to obtain soluble fractions, and the remaining pellets were further extracted to recover insoluble Aβ. Soluble Aβ was extracted in TBS or PBS-based buffer, and insoluble Aβ was extracted from pellets using 5 M guanidine-HCl or 70% formic acid. Aβ42 levels were quantified using a commercially available ELISA kit (Thermo Fisher Scientific; Aβ42, Cat. # KMB3441), according to the manufacturer’s instructions. Concentrations were interpolated from standard curves and normalized to total protein content determined by BCA assay. Data are reported separately for cortex and hippocampus.

### Cytokine analysis

Brain homogenates were prepared in lysis buffer and cytokine levels (IL-1β, TNF-α) were measured using ELISA kits (R&D Systems, catalog# MLB00C and MTA00B, respectively). Values were normalized to total protein content.

### Microglial activation

Brain tissues (cortex and hippocampus) were rapidly dissected and processed into single-cell suspensions using the Adult Brain Dissociation Kit (Miltenyi Biotec, Cat. #130-107-677) according to the manufacturer’s protocol. Myelin was removed using Myelin Removal Beads II (Miltenyi Biotec, Cat. #130-096-733). Cells were resuspended in FACS buffer (PBS containing 2% fetal bovine serum and 2 mM EDTA) and incubated with anti-mouse CD16/CD32 Fc block (BioLegend, Cat. #101320) for 10 min at 4 °C to prevent non-specific binding.

Cells were stained for 30 min at 4 °C with fluorophore-conjugated antibodies against CD45 (APC, clone 30-F11, BioLegend, Cat. #103112), CD11b (FITC, clone M1/70, BioLegend, Cat. #101206), TMEM119 (PE, clone 106-6, BioLegend, Cat. #157306), and CD86 (PerCP-Cy5.5, clone GL-1, BioLegend, Cat. #105028). Dead cells were excluded using a viability dye (Zombie NIR, BioLegend, Cat. #423106). Data were acquired on a BD LSRFortessa flow cytometer (BD Biosciences) and analyzed using FlowJo v10. Microglia were defined as live CD45^low^ CD11b^+^ TMEM119^+^ cells. Activated microglia were quantified as the percentage of CD86^+^ cells within the microglial population. A minimum of 50,000 live events per sample were recorded.

### Statistical analysis

All data are presented as mean ± SD. Statistical analyses were performed using GraphPad Prism v10. Two-group comparisons were conducted using unpaired two-tailed Student’s *t*-tests. Multiple comparisons were analyzed using one-way ANOVA followed by Tukey’s post hoc test. *P* values < 0.05 were considered statistically significant. Exact sample sizes and statistical tests are indicated in figure legends.

## ASSOCIATED CONTENT

The following files are available free of charge.

Top-ranked compounds from AI-based screening, LILRB4 binding affinities, ELISA and BLI validation studies, and cross species LILRB4 reactivity (PDF).

## Notes

The authors declare no competing financial interest.

---

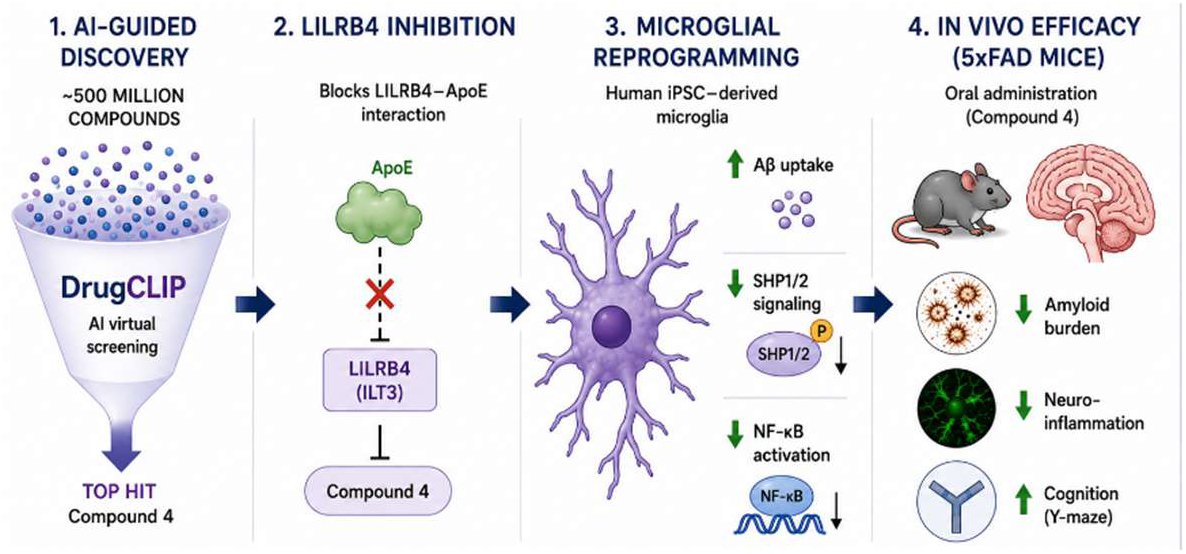

