## Supporting Information for "AI-Guided Discovery of Small Molecule LILRB4 (ILT3) Inhibitors Reprograms Microglia and Reduces Amyloid Pathology"

*Electronic Supplementary Information for*

| **Contents** |  |
| --- | --- |
| Top-ranked compounds from DrugCLIP screening  Binding affinities of validated hits as LILRB4 binders measured by MST  Dose-dependent inhibition of ApoE binding to LILRB4 by compound **15** measured by ELISA | S3  S8  S8 |
| BLI-based measurement of compound **15** activity for LILRB4-ApoE inhibition showing a concentration-dependent decrease in binding response | S9 |
| Cross-species binding of compound **4** to murine LILRB4 measured by MST | S10 |

**Table S1. Top-ranked compounds from DrugCLIP screening.** Candidates are listed with molecule ID, source library, SMILES, DrugCLIP score, and AutoDock Vina docking score (kcal·mol⁻¹). Compounds were prioritized based on combined DrugCLIP ranking and predicted binding affinity.

| **Comp. No.** | **Mol ID** | **Library** | **SMILES** | **DrugCLIP Score** | **Vina Score** |
| --- | --- | --- | --- | --- | --- |
| **1** | OSSL_419185 | Princeton_BioMolecular_Research | c1ccccc1-n(n(C)c2C)c(=O)c2NC(=O)CN(S(=O)(=O)C)c(cc3)ccc3C45CC6CC(C5)CC(C4)C6 | 2.88 | -6.78 |
| **2** | STL026261 | Vitas-M | c1ccccc1-n(n(C)c2C)c(=O)c2NC(=O)CN(S(=O)(=O)C)c(cc3)ccc3C45CC6CC(C5)CC(C4)C6 | 2.86 | -6.51 |
| **3** | C201-0799 | ChemDiv | CC(C)Cc(cc1)ccc1S(=O)(=O)c2c(=O)n(c(C)cc2C)CC(=O)Nc(cc3)ccc3C(=O)OCC | 2.84 | -6.13 |
| **4** | V010-0630 | ChemDiv | FC(F)(F)c1cc(ccc1)-c(on2)nc2-c3ccc(cc3)CN(CC4)CCN4C(=O)C(NC(=O)C)CC(C)C | 2.8 | -7.85 |
| **5** | OSSL_470920 | Princeton_BioMolecular_Research | COC(=O)c1ccc(cc1)\C=N\NC(=O)CN(S(=O)(=O)C)c(cc2)ccc2C34CC5CC(C3)CC(C4)C5 | 2.73 | -6.29 |
| **6** | STL090215 | Vitas-M | COC(=O)c1ccc(cc1)\C=N\NC(=O)CN(S(=O)(=O)C)c(cc2)ccc2C34CC5CC(C3)CC(C4)C5 | 2.72 | -6.35 |
| **7** | STL038215 | Vitas-M | c1ccccc1-n2c(-c3ccc(C(C)(C)C)cc3)n[nH+]c2SCC(=O)N/N=C/c(c4[O-])cccc4CC=C | 2.72 | -6.52 |
| **8** | STK071041 | Vitas-M | c1cc(C)c(C)cc1N(S(=O)(=O)C)Cc(cc2)ccc2C(=O)N/N=C/c(c3)ccc(c3OCC)OCC=C | 2.72 | -6.17 |
| **9** | OSSK_368196 | Princeton_BioMolecular_Research | c1cc(C)c(C)cc1N(S(=O)(=O)C)Cc(cc2)ccc2C(=O)N/N=C/c(c3)ccc(c3OCC)OCC=C | 2.71 | -5.87 |
| **10** | OSSL_455296 | Princeton_BioMolecular_Research | COCCNC(=O)c1c(cccc1)NC(=O)CN(S(=O)(=O)C)c(cc2)ccc2C34CC5CC(C4)CC(C3)C5 | 2.7 | -6.31 |
| **11** | Z4898060549 | Enamine_screening_collection_sdf_202504 | COC(=O)c1cccc(c1)C#Cc2cccc(c2)NC(=O)C(=O)N(C)C3CCCC3c4ccc(C)cc4 | 2.7 | -6.87 |
| **12** | V009-0312 | ChemDiv | CCCc(cc1)ccc1S(=O)(=O)N2CCN(CC2)Cc(oc3)cc(=O)c3OCc(cc4)ccc4C(C)(C)C | 2.69 | -6.28 |
| **13** | STL090217 | Vitas-M | COc(c1)ccc(c1OC)\C=N\NC(=O)CN(S(=O)(=O)C)c(cc2)ccc2C34CC5CC(C4)CC(C3)C5 | 2.68 | -6.38 |
| **14** | V007-0166 | ChemDiv | c1ccc(F)cc1-n2nc(C(=O)OCC)cc2-c3ccc(cc3)N(CC4)CCN4C(=O)C(C)Oc5ccccc5 | 2.68 | -6.84 |
| **15** | STK316238 | Vitas-M | CC(C)c1ccc(cc1)OCc(c2)cccc2C(=O)Nc(ccc3)cc3S(=O)(=O)N4CCOCC4 | 2.68 | -7.02 |
| **16** | E823-0045 | ChemDiv | CC(C)Cc(cc1)ccc1S(=O)(=O)N2CCN(CC2)Cc3nc(on3)CCC(=O)N(C)C4CCCCC4 | 2.67 | -5.59 |
| **17** | STK190354 | Vitas-M | c1ccccc1C(O)(c2ccccc2)C(=O)N/N=C/c(cc3)ccc3OC(=O)/C=C/c4ccc(cc4)OC | 2.67 | -6.05 |
| **18** | OSSK_615206 | Princeton_BioMolecular_Research | c1ccccc1C(O)(c2ccccc2)C(=O)N/N=C/c(cc3)cc(OCC)c3OC(=O)c4ccc(C)cc4 | 2.67 | -6.29 |
| **19** | STL090214 | Vitas-M | COc1c(OC)ccc(c1)\C=N\NC(=O)CN(S(=O)(=O)C)c(cc2)ccc2C34CC5CC(C4)CC(C3)C5 | 2.66 | -6.07 |
| **20** | Y501-3938 | ChemDiv | C1COCCN1S(=O)(=O)c2cc(ccc2)NC(=O)c3cc(ccc3)COc(cc4)ccc4C(C)C | 2.64 | -7.21 |
| **21** | V007-0120 | ChemDiv | c1cc(F)ccc1-n2nc(C(=O)OCC)cc2-c3ccc(cc3)N(CC4)CCN4C(=O)C(C)Oc5ccccc5 | 2.64 | -6.66 |
| **22** | Z31026090 | Enamine_screening_collection_sdf_202504 | CCc1ccc(cc1)NC(=O)c2ccccc2NCC(=O)Nc3cc(ccc3C)S(=O)(=O)N4CCOCC4 | 2.64 | -7.01 |
| **23** | V012-9558 | ChemDiv | C1CCCC1c(on2)nc2-c3ccc(cc3)CN(CC4)CCN4C(=O)C(NC(=O)C)Cc5c(F)cccc5 | 2.64 | -6.49 |
| **24** | OSSL_470922 | Princeton_BioMolecular_Research | COc(c1)ccc(c1OC)\C=N\NC(=O)CN(S(=O)(=O)C)c(cc2)ccc2C34CC5CC(C4)CC(C3)C5 | 2.64 | -6 |
| **25** | V009-7855 | ChemDiv | c1cccc(C)c1-n2nc(C(=O)OCC)cc2-c3ccc(cc3)N(CC4)CCN4C(=O)COc5ccccc5 | 2.63 | -6.12 |
| **26** | STK190383 | Vitas-M | c1ccccc1C(O)(c2ccccc2)C(=O)N/N=C/c(cc3)cc(OCC)c3OC(=O)c4ccc(C)cc4 | 2.63 | -6.19 |
| **27** | STL032582 | Vitas-M | COCCNC(=O)c1c(cccc1)NC(=O)CN(S(=O)(=O)C)c(cc2)ccc2C34CC5CC(C4)CC(C3)C5 | 2.62 | -6.08 |
| **28** | OSSL_922481 | Princeton_BioMolecular_Research | c1ccc(C)c(C)c1N(CC2)CCN2S(=O)(=O)c3c(cn(n3)CC)C(=O)NCc4ccc(cc4)OCCC | 2.62 | -6.44 |
| **29** | Z90098581 | Enamine_screening_collection_sdf_202504 | CC(C)(C)c1ccc(cc1)NC(=O)CSc2ccc(cn2)S(=O)(=O)N3CCN(CC3)Cc4ccccc4 | 2.62 | -5.49 |
| **30** | STL090213 | Vitas-M | COc(c1)ccc(OC)c1\C=N\NC(=O)CN(S(=O)(=O)C)c(cc2)ccc2C34CC5CC(C4)CC(C3)C5 | 2.62 | -6.23 |
| **31** | OSSL_470919 | Princeton_BioMolecular_Research | COc1c(OC)ccc(c1)\C=N\NC(=O)CN(S(=O)(=O)C)c(cc2)ccc2C34CC5CC(C4)CC(C3)C5 | 2.62 | -5.7 |
| **32** | V004-6494 | ChemDiv | c1ccccc1C(OC(=O)C)C(=O)N2CCN(CC2)Cc(oc3)cc(=O)c3OCc(cc4)ccc4C(C)(C)C | 2.62 | -6.64 |
| **33** | K061-0077 | ChemDiv | c1cc(OC)ccc1-c(cc2C(F)(F)F)nc(n23)cc(n3)C(=O)NC(C(=O)OC)C45CC6CC(C5)CC(C4)C6 | 2.61 | -6.86 |
| **34** | Z106310838 | Enamine_screening_collection_sdf_202504 | Cc1cccc(c1C)NC(=O)CN2CCN(CC2)CC(=O)NC(C(C)C)c3ccc(cc3)C(C)C | 2.61 | -7 |
| **35** | D516-0148 | ChemDiv | C1CCCC1C(=O)NCCc(n2)n(C)c(c23)ccc(c3)NC(=O)COc(cc4)ccc4C(C)(C)C | 2.61 | -6.75 |
| **36** | Z197773058 | Enamine_screening_collection_sdf_202504 | Cc1ccc(cc1C)-c2nn(-c3ccccc3)cc2C(=O)NCc4ccc(cc4)CS(=O)(=O)NC(C)C | 2.61 | -6.81 |
| **37** | OSSL_470918 | Princeton_BioMolecular_Research | COc(c1)ccc(OC)c1\C=N\NC(=O)CN(S(=O)(=O)C)c(cc2)ccc2C34CC5CC(C4)CC(C3)C5 | 2.61 | -6.48 |
| **38** | V006-0654 | ChemDiv | c1cccc(C)c1-n2nc(C(=O)OCC)cc2-c3ccc(cc3)N(CC4)CCN4C(=O)Cc5ccc(cc5)OC | 2.6 | -6.37 |
| **39** | OSSK_616667 | Princeton_BioMolecular_Research | CCCCC(=O)Nc1c(OC)cc(cc1)NCc(c2)ccc(c2OC)OCC(=O)NC(C)(C)C | 2.6 | -5.91 |
| **40** | Z51812895 | Enamine_screening_collection_sdf_202504 | c1ccccc1-n(n2C)c(=O)c(c2C)NC(=O)c3cccc(c3)NC(=O)Cc(cc4)cc(c45)CCCC5 | 2.6 | -7.75 |
| **41** | STL133115 | Vitas-M | c1ccc(C)c(C)c1N(CC2)CCN2S(=O)(=O)c3c(cn(n3)CC)C(=O)NCc4ccc(cc4)OCCC | 2.6 |  |
| **42** | Z56060483 | Enamine_screening_collection_sdf_202504 | c1ccccc1-n(n2C)c(=O)c(c2C)NC(=O)c3cccc(c3)NC(=S)NC(=O)C45CC6CC(C5)CC(C6)C4 | 2.6 | -7.14 |
| **43** | G740-1338 | ChemDiv | c1ccccc1-n(c2C)nc(c23)c(C)nn(c3=O)C(CC)C(=O)NCc4ccc(cc4)OCCCC | 2.59 | -5.44 |
| **44** | F195-0764 | ChemDiv | CC(C)Cc(cc1)ccc1S(=O)(=O)c2c(=O)n(c(C)cc2C)CC(=O)Nc3cc(OC)cc(c3)OC | 2.59 | -6.8 |
| **45** | Z216985454 | Enamine_screening_collection_sdf_202504 | CC(C)COc1ccc(cc1OC)/C=C/C(=O)NCc2ccccc2S(=O)(=O)N3CCOCC3 | 2.59 | -5.76 |
| **46** | Z4898032576 | Enamine_screening_collection_sdf_202504 | Cc1cccc(c1)COc2ccc(cc2)C(C)NC(=O)Nc3cccc(c3)S(=O)(=O)N4CCN(C)CC4 | 2.58 | -6.56 |
| **47** | Z220070482 | Enamine_screening_collection_sdf_202504 | Cc1ccc(cc1)NC(=O)CN2CCN(CC2)CCC(=O)Nc3ccccc3Oc4ccccc4 | 2.58 | -6.06 |
| **48** | V006-0290 | ChemDiv | c1cc(F)ccc1-n2nc(C(=O)OCC)cc2-c3ccc(cc3)N(CC4)CCN4C(=O)Cc5ccc(F)cc5 | 2.58 | -6.75 |
| **49** | STK424251 | Vitas-M | C1C=CCC(C(=O)O)C1C(=O)Nc(cc(c23)OCO3)c(c2)/C(C)=N/NC(=O)CC45CC6CC(C5)CC(C4)C6 | 2.58 | -7.13 |
| **50** | Z96242015 | Enamine_screening_collection_sdf_202504 | C1COCCN1S(=O)(=O)c(ccc2OCC)cc2NC(=O)COc3ccc(cc3)C(C)(C)CC | 2.58 | -5.75 |
| **51** | OSSK_615173 | Princeton_BioMolecular_Research | c1ccccc1C(O)(c2ccccc2)C(=O)N/N=C/c(cc3)ccc3OC(=O)/C=C/c4ccc(cc4)OC | 2.58 | -6.44 |
| **52** | V008-0238 | ChemDiv | c1ccc(F)cc1C(=O)N(CC=C)CC(=O)N2CCN(CC2)c3nnc(cc3)-c4ccc(cc4)-c5ccccc5 | 2.58 | -6.3 |
| **53** | OSSL_028508 | Princeton_BioMolecular_Research | C1COCCN1S(=O)(=O)c2cc(ccc2)NC(=O)c3cc(ccc3)COc(cc4)ccc4C(C)C | 2.57 | -7.06 |
| **54** | Z30376622 | Enamine_screening_collection_sdf_202504 | C1COCCN1S(=O)(=O)c(ccc2OC(C)C)cc2NC(=O)COc3ccc(cc3)C(C)(C)C | 2.56 | -6.13 |
| **55** | V003-1639 | ChemDiv | CC(C)Cc(on1)nc1-c2ccc(cc2)CN(CC3)CCN3C(=O)C(NC(=O)C)Cc4c(F)cccc4 | 2.56 | -6.95 |
| **56** | OSSL_922474 | Princeton_BioMolecular_Research | CC(C)c1ccc(cc1)CNC(=O)c(cn(n2)CC)c2S(=O)(=O)N3CCN(CC3)c4c(C)c(C)ccc4 | 2.56 | -6.91 |
| **57** | F195-0767 | ChemDiv | CC(C)Cc(cc1)ccc1S(=O)(=O)c2c(=O)n(c(C)cc2C)CC(=O)Nc3ccc(cc3)OCC | 2.56 | -6.52 |
| **58** | Z3271810801 | Enamine_screening_collection_sdf_202504 | CCCCc1ccc(cc1)NS(=O)(=O)c2ccc(C)c(c2)C(=O)Nc3cccc(c3)CN4CCCC4C(=O)N | 2.56 | -6.87 |
| **59** | OSSL_470849 | Princeton_BioMolecular_Research | c1ccccc1N(S(=O)(=O)C)Cc(cc2)ccc2C(=O)Nc(cccc3)c3C(=O)NCCc4ccccc4 | 2.56 | -6.58 |
| **60** | STL038182 | Vitas-M | c1cc(C)ccc1-n2c(-c3ccc(C(C)(C)C)cc3)n[nH+]c2SCC(=O)N/N=C/c(c4[O-])cccc4OCC | 2.56 | -7.04 |
| **61** | Z44839698 | Enamine_screening_collection_sdf_202504 | c1cc(C)ccc1S(=O)(=O)Oc2ccc(cc2OC)/C=N/NC(=O)COc3ccc(cc3)C(C)(C)C | 2.55 | -6.09 |
| **62** | E823-0150 | ChemDiv | CC(C)Cc(cc1)ccc1S(=O)(=O)N2CCN(CC2)Cc3nc(on3)CCC(=O)N4CCCCCC4 | 2.55 | -6.1 |
| **63** | V003-0139 | ChemDiv | c1ccc(F)cc1-n2nc(C(=O)OCC)cc2-c3ccc(cc3)N(CC4)CCN4C(=O)Cc5ccc(cc5)OC | 2.55 | -6.61 |
| **64** | Z4416716924 | Enamine_screening_collection_sdf_202504 | CC(F)(F)c1ccc(nc1)Cn2cc(nn2)COc3ccccc3CNC(=O)C45CCC(CC4)(CC5)C(=O)O | 2.55 | -6.98 |
| **65** | V008-0302 | ChemDiv | c1ccccc1C(=O)N(CC=C)CC(=O)N2CCN(CC2)c3nnc(cc3)-c4ccc(cc4)-c5ccccc5 | 2.55 | -6.4 |
| **66** | V014-8990 | ChemDiv | CCC(CC)c1cc(C(=O)OCC)nn1-c2ccc(cc2)C(=O)NCCc(c3)ccc(OC)c3OC | 2.54 | -6.34 |
| **67** | D073-0208 | ChemDiv | c1ccccc1C(=O)NCCc(n2)n(C)c(c23)ccc(c3)NC(=O)COc4c(C(C)C)ccc(c4)C | 2.54 | -6 |
| **68** | Z200681008 | Enamine_screening_collection_sdf_202504 | Cc1ccc(cc1)CCC2CCN(CC2)C(=O)CCc(n3)n(C)c(c34)ccc(c4)S(=O)(=O)N5CCOCC5 | 2.54 | -6.91 |
| **69** | STK165952 | Vitas-M | c1ccccc1-c(nn2)nn2CC(=O)N/N=C/c(cc3)ccc3OC(=O)c4ccc(C(C)(C)C)cc4 | 2.54 | -6.75 |
| **70** | E823-0060 | ChemDiv | CC(C)Cc(cc1)ccc1S(=O)(=O)N2CCN(CC2)Cc3nc(on3)CCC(=O)N4CCC(C)CC4 | 2.54 | -6.61 |
| **71** | Z9296883777 | Enamine_screening_collection_sdf_202504 | C/C=C/COc1cccc(c1)CC(=O)NCc2cn(nn2)C3CN(C3)c4oc(C(C)(C)C)nc4C#N | 2.54 | -6.19 |
| **72** | V006-0310 | ChemDiv | c1ccc(F)cc1-n2nc(C(=O)OCC)cc2-c3ccc(cc3)N(CC4)CCN4C(=O)Cc5ccc(F)cc5 | 2.54 | -7.34 |
| **73** | L861-0679 | ChemDiv | c1cc(C)ccc1CNC(=O)c(c2)ccc(n23)nn(c3=O)CC(=O)Nc4ccc(cc4)CCCC | 2.54 | -6.82 |
| **74** | STK499770 | Vitas-M | c1ccccc1C(=O)NCCc(n2)n(C)c(c23)ccc(c3)NC(=O)COc4c(C(C)C)ccc(c4)C | 2.53 | -6.28 |
| **75** | Z285604762 | Enamine_screening_collection_sdf_202504 | C1CCCCN1S(=O)(=O)c(ccc2C)cc2NC(=O)CNc3cccc(c3)C#Cc4ccccc4 | 2.53 | -6.89 |
| **76** | STL376075 | Vitas-M | CC(C)OC(=O)CN1C(=O)SC(\C1=O)=C\c2ccc(cc2)OCC(=O)Nc3c(C(C)C)cccc3 | 2.53 | -6.98 |
| **77** | D647-0123 | ChemDiv | CCOc(cc1)ccc1-c(c2)onc2C(=O)NCC(N3CCOCC3)c(cc4)ccc4C(C)(C)C | 2.53 | -6.23 |
| **78** | V015-1302 | ChemDiv | c1cc(OC)ccc1-n2nc(C(=O)OCC)cc2-c3ccc(cc3)N(CC4)CCN4C(=O)Cc5ccccc5 | 2.53 | -6.29 |
| **79** | K940-1914 | ChemDiv | CCOc(cc1)ccc1NC(=O)CN(CC2)CCN2CC(=O)Nc3ccc(cc3)OCc4ccccc4 | 2.53 | -6.23 |
| **80** | V007-0108 | ChemDiv | c1cccc(C)c1-n2nc(C(=O)OCC)cc2-c3ccc(cc3)N(CC4)CCN4C(=O)Cc5ccccc5 | 2.52 | -6.35 |
| **81** | V008-7522 | ChemDiv | c1ccc(F)cc1-n2nc(C(=O)OCC)cc2-c3ccc(cc3)N(CC4)CCN4C(=O)Cc5cc(OC)ccc5 | 2.52 | -6.65 |
| **82** | OSSL_675337 | Princeton_BioMolecular_Research | C1COCCN1C(=O)C2C(c3ccccc3)C(C2c4ccccc4)C(=O)N/N=C/c(cc5)ccc5N(CC)CC | 2.52 | -6.36 |
| **83** | V002-9936 | ChemDiv | c1cc(F)ccc1-n2nc(C(=O)OCC)cc2-c3ccc(cc3)N(CC4)CCN4C(=O)Cc5ccc(cc5)OC | 2.52 | -6.76 |
| **84** | STL362416 | Vitas-M | c1ccccc1-n2c(-c3ccc(C(C)(C)C)cc3)nnc2SCC(=O)N/N=C/c4c(OCC(=O)O)cccc4 | 2.52 | -6.8 |
| **85** | OSSL_258837 | Princeton_BioMolecular_Research | c1ccccc1C(=O)NCCc(n2)n(C)c(c23)ccc(c3)NC(=O)COc4c(C(C)C)ccc(c4)C | 2.51 | -6.99 |
| **86** | OSSK_537175 | Princeton_BioMolecular_Research | c1ccccc1-c(nn2)nn2CC(=O)N/N=C/c(cc3)ccc3OC(=O)c4ccc(C(C)(C)C)cc4 | 2.51 | -6.42 |
| **87** | D516-0346 | ChemDiv | C1CCCC1C(=O)NCCc(n2)n(C)c(c23)ccc(c3)NC(=O)COc4c(C(C)C)ccc(c4)C | 2.51 | -6.06 |
| **88** | Z437269656 | Enamine_screening_collection_sdf_202504 | c1cc(C)ccc1C(NC(=O)C)CC(=O)N2CCN(CC2)C(=O)c3ccc(cc3)OCc4ccccc4 | 2.51 | -6.95 |
| **89** | STK613148 | Vitas-M | CC(C)COc(cc1)ccc1C(=O)N/N=C(C)/C=N/NC(=O)c2ccc(cc2)OCC(C)C | 2.51 | -5.43 |
| **90** | OSSL_109315 | Princeton_BioMolecular_Research | CCOc(cc1)ccc1-c(c2)onc2C(=O)NCC(N3CCOCC3)c(cc4)ccc4C(C)(C)C | 2.5 | -5.78 |
| **91** | V023-5801 | ChemDiv | c1cc(C)ccc1Cn2c(=O)n(-c3ccc(cc3)C(C)C)nc(c2=O)C(=O)N4CC(CCC4)C(=O)OCC | 2.5 | -6.92 |
| **92** | Z25794881 | Enamine_screening_collection_sdf_202504 | C1CCCCCC1NC(=O)CO\N=C\c(cc2OC)ccc2OCC(=O)Nc3c(C)cc(C)cc3C | 2.5 | -6.62 |
| **93** | Z31731818 | Enamine_screening_collection_sdf_202504 | c1ccccc1-n(n2C)c(=O)c(c2C)NS(=O)(=O)c3cccc(c3)C(=O)Nc4ccc(cc4)C(C)(C)C | 2.5 | -6.55 |
| **94** | Z56796886 | Enamine_screening_collection_sdf_202504 | CC(C)(C)c1ccc(cc1)-c2nnc(n2-c3ccccc3)SCC(=O)N/N=C/c4ccccc4OCC(=O)O | 2.5 | -6.39 |
| **95** | E823-0037 | ChemDiv | c1cc(C(C)(C)C)ccc1S(=O)(=O)N2CCN(CC2)Cc3nc(on3)CCC(=O)N(C)C4CCCCC4 | 2.5 | -5.94 |
| **96** | OSSL_149586 | Princeton_BioMolecular_Research | c1cccc(OCC)c1NC(=O)C(=O)N/N=C/c2c(cccc2)OCC(=O)Nc(cc3C(F)(F)F)ccc3 | 2.5 | -6.88 |
| **97** | STK470227 | Vitas-M | c1ccccc1C(O)(c2ccccc2)C(=O)N/N=C(C)/c3ccc(cc3)NC(=O)C4CCCCC4 | 2.5 | -6.1 |
| **98** | Z26762628 | Enamine_screening_collection_sdf_202504 | C1COCCN1S(=O)(=O)c(c2)ccc(c23)n(CCC)c(n3)CCC(=O)Nc4ccccc4Cc5ccccc5 | 2.5 | -6.64 |
| **99** | STL350237 | Vitas-M | c1ncnn1Cc(cc2)ccc2NC(=O)Cc(c3C)c(=O)oc(c4)c3cc(CC5)c4OC56CCCCC6 | 2.49 | 72.85 |
| **100** | STK977689 | Vitas-M | c1ccccc1N(S(=O)(=O)C)Cc(cc2)ccc2C(=O)Nc(cccc3)c3C(=O)NCCc4ccccc4 | 2.49 | -6.85 |

**Table 2. Binding affinities of validated hits as LILRB4 binders measured by MST.**
Equilibrium dissociation constants (Kd, µM) for the compounds were determined using microscale thermophoresis (MST).

| **Comp. No.** | **MST KD (μM)** |
| --- | --- |
| **4** | 0.02 |
| **15** | 0.09 |
| **56** | 0.45 |
| **32** | 2.63 |
| **20** | 8.51 |
| **78** | 12.4 |
| **55** | 19.5 |
| **66** | 22.5 |
| **49** | 34.2 |

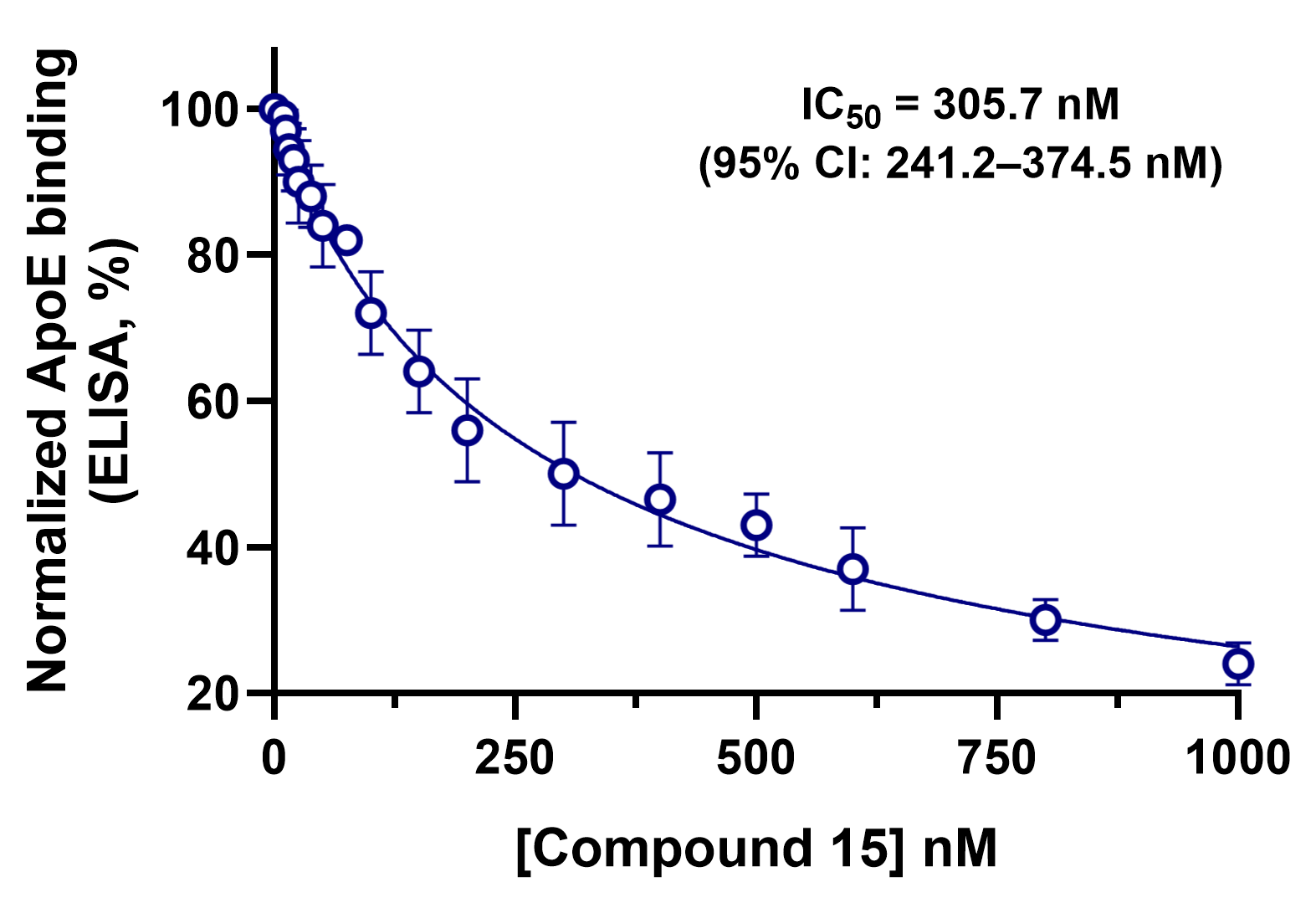

**Figure S1.** Dose-dependent inhibition of ApoE binding to LILRB4 by compound **15** measured by ELISA. Data are presented as normalized ApoE binding (%). Data are shown as mean ± SD (n = 5).

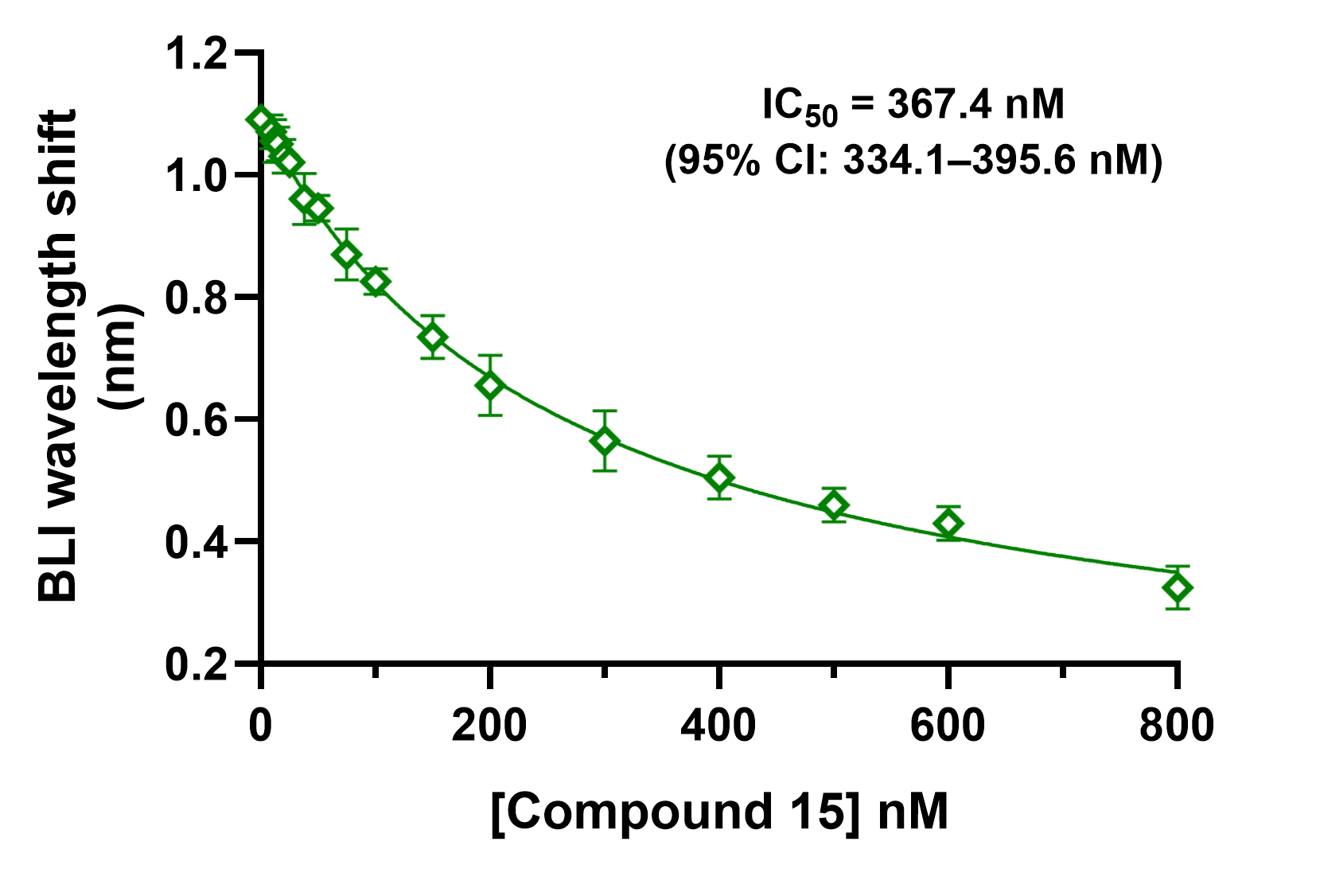

**Figure S2.** BLI-based measurement of compound **15** activity for LILRB4-ApoE inhibition showing a concentration-dependent decrease in binding response. Data are shown as mean ± SD (n = 5).

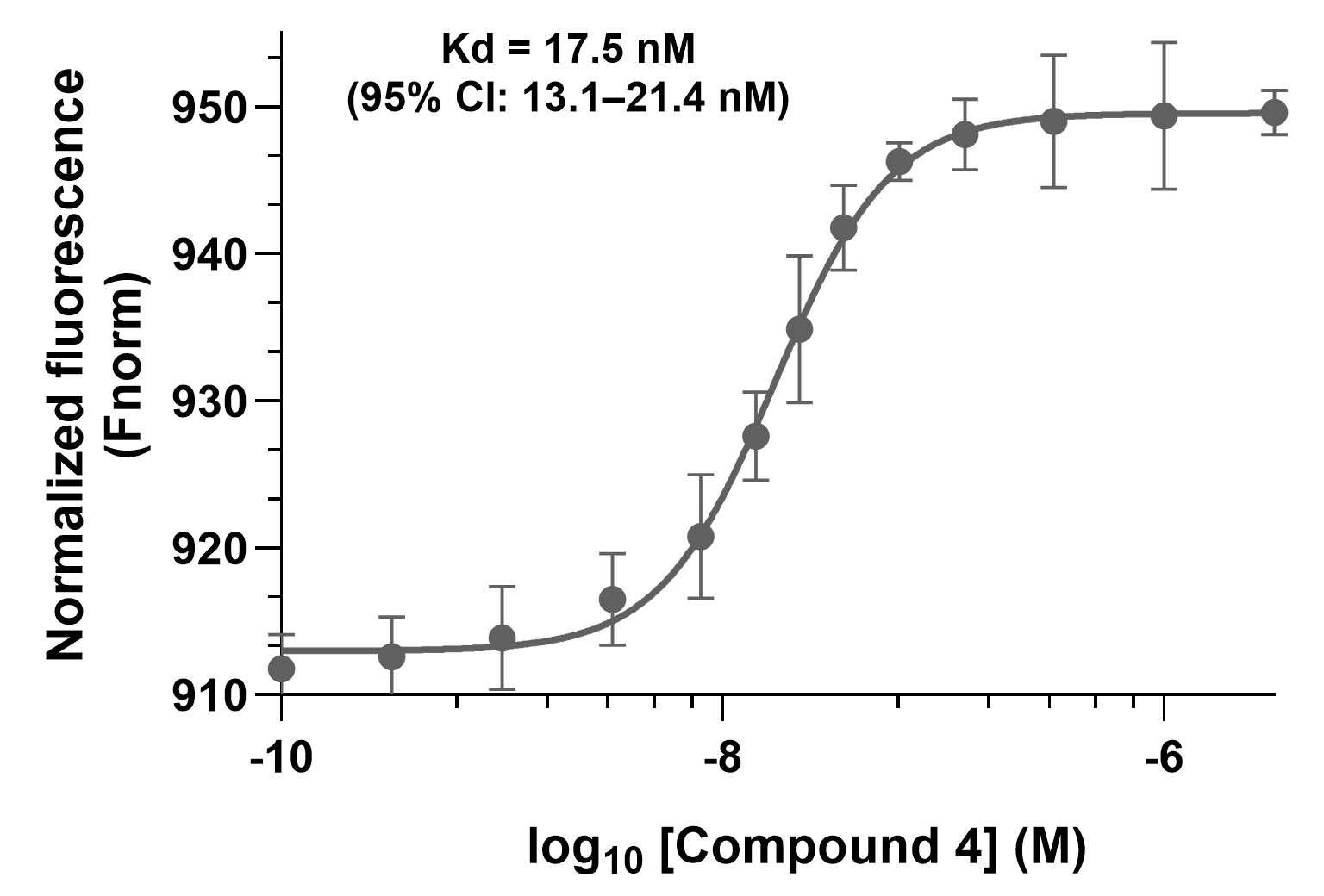

**Figure S3.** **Cross-species binding of compound 4 to murine LILRB4 measured by MST.** Normalized thermophoresis signals were recorded across a concentration series of compound **4** against the recombinant extracellular domain of mouse LILRB4. Data shows a concentration-dependent change in thermophoretic response consistent with specific binding. Data are presented as mean ± SD (n = 5).
